# Evaluation of whole genome amplification and bioinformatic methods for the characterization of *Leishmania* genomes at a single cell level

**DOI:** 10.1101/2020.02.20.957621

**Authors:** Hideo Imamura, Marlene Jara, Pieter Monsieurs, Mandy Sanders, Ilse Maes, Manu Vanaerschot, Matthew Berriman, James A. Cotton, Jean-Claude Dujardin, Malgorzata A. Domagalska

## Abstract

Here, we report a pilot study paving the way for further single cell genomics studies in *Leishmania*. First, the performances of two commercially available kits for Whole Genome Amplification (WGA), PicoPlex and RepliG was compared on small amounts of *Leishmania donovani* DNA, testing their ability to preserve specific genetic variations, including aneuploidy levels and SNPs. We show here that the choice of WGA method should be determined by the planned downstream genetic analysis, Picoplex and RepliG performing better for aneuploidy and SNP calling, respectively. This comparison allowed us to evaluate and optimize corresponding bio-informatic methods. As PicoPlex was shown to be the preferred method for studying single cell aneuploidy, this method was applied in a second step, on single cells of *L. braziliensis*, which were sorted by fluorescence activated cell sorting (FACS). Even sequencing depth was achieved in 28 single cells, allowing accurate somy estimation. A dominant karyotype with three aneuploid chromosomes was observed in 25 cells, while two different minor karyotypes were observed in the other cells. Our method thus allowed the detection of aneuploidy mosaicism, and provides a solid basis which can be further refined to concur with higher-throughput single cell genomic methods.

## Introduction

*Leishmania* are unicellular protozoan parasites belonging to the family *Trypanosomatidae* and causing a spectrum of diseases in tropical and sub-tropical regions, with an incidence estimated at 1.6 million cases per year ^1^. The parasite has a dimorphic life cycle: extracellular flagellated promastigotes in the sand fly vector and intracellular amastigotes in macrophages of the vertebrate host. *Leishmania* have unique genetic and molecular properties that distinguish them from other unicellular and multicellular eukaryotes. Among others, there is no transcription regulation at initiation through promoters. Instead, genes are organized in long arrays of polycistronic units that are transcribed together, then trans-spliced and polyadenylated, and their expression is regulated post-transcriptionally^2^. In this context, gene dosage represents a straightforward strategy for the parasite to modify the expression of genes of interest ^3^. This can occur through different mechanisms, like expansion/contraction of tandem arrays, episomal amplifications and aneuploidy^4^.

We previously sequenced the genomes of 204 cultivated clinical isolates of *L. donovani* and found aneuploidy in all of them, often affecting half of the 36 chromosomes^5^. Experimental evolution studies suggest that aneuploidy changes constitute an adaptive mechanism to environmental changes like those occurring during the life cycle^6^ or those associated with drug pressure^7^. In contrast to many organisms where it can be deleterious, aneuploidy thus appears to be crucial in *Leishmania*. Analyses of aneuploidy at individual promastigote cell level by FISH revealed an additional dimension, i.e. the concept of mosaic aneuploidy: the tested chromosomes were present in two or more somy states (from monosomic to tetrasomic), varying between cells in a clonal line, so that the karyotype varies from cell to cell^8^. Mosaic aneuploidy could originate from segregation defects, but the currently favored hypothesis is a defect in regulation of chromosome replication^9^.

So far, mosaic aneuploidy was only studied for a few *Leishmania* chromosomes, and these pioneering FISH-based studies should be complemented and refined by single cell genomic methods. A range of different methods exist, in which cell- or nucleus-sorting is coupled with whole genome amplification (WGA) and high-throughput sequencing^10^. Single cell genomics is well established in human genetics and cancer research among others^11^, but it constitutes an emerging field in parasitology, with a few published studies in parasites like *Plasmodium*^12^ or *Cryptosporidium*^13^, never -to our knowledge- in *Leishmania* and other Trypanosomatids. We report here the first study on single cell genomics in *Leishmania*. In this pilot study, we first compared the performances of two WGA methods on their ability to preserve specific genetic variations, including aneuploidy levels and SNPs, using small amounts of *Leishmania* DNA. Alongside the assessment of both methods to detect aneuploidy and SNPs at the single cell level, this comparison allowed us to evaluate and optimize corresponding bioinformatic methods. The PicoPlex approach was more adequate to assess chromosome number and we applied it to study aneuploidy of single cells sorted by fluorescence activated cell sorting (FACS). This allowed us to distinguish the dominating karyotype from minor ones at the single cell level.

## Results

### Genome coverage and base accuracy

In a first part of the study, we used DNA extracted from different numbers of cells (from 10 to 1000) of *L. donovani* BPK275 to compare the performance of RepliG and PicoPlex WGA methods and optimize the downstream bioinformatic protocols. The average depth for RepliG was significantly higher than that of PicoPlex. Specifically, for the five RepliG samples, the total number of reads was 222 million and 96.2 % of the reads were mapped to the *L. donovani* BPK282 reference, resulting in average depth of 17.1 ± 14.7. For the five PicoPlex samples, the total number of reads was 111 million and 41.1 % of the reads were mapped, resulting in average depth 3.4 ± 2.4. (Supplementary data 1, Table S1). As somy calculation for RepliG_10 sample was not successful due to uneven sequencing depth resulting in chromosomes with almost no coverage, this sample was omitted in downstream processing.

**Table 1.**
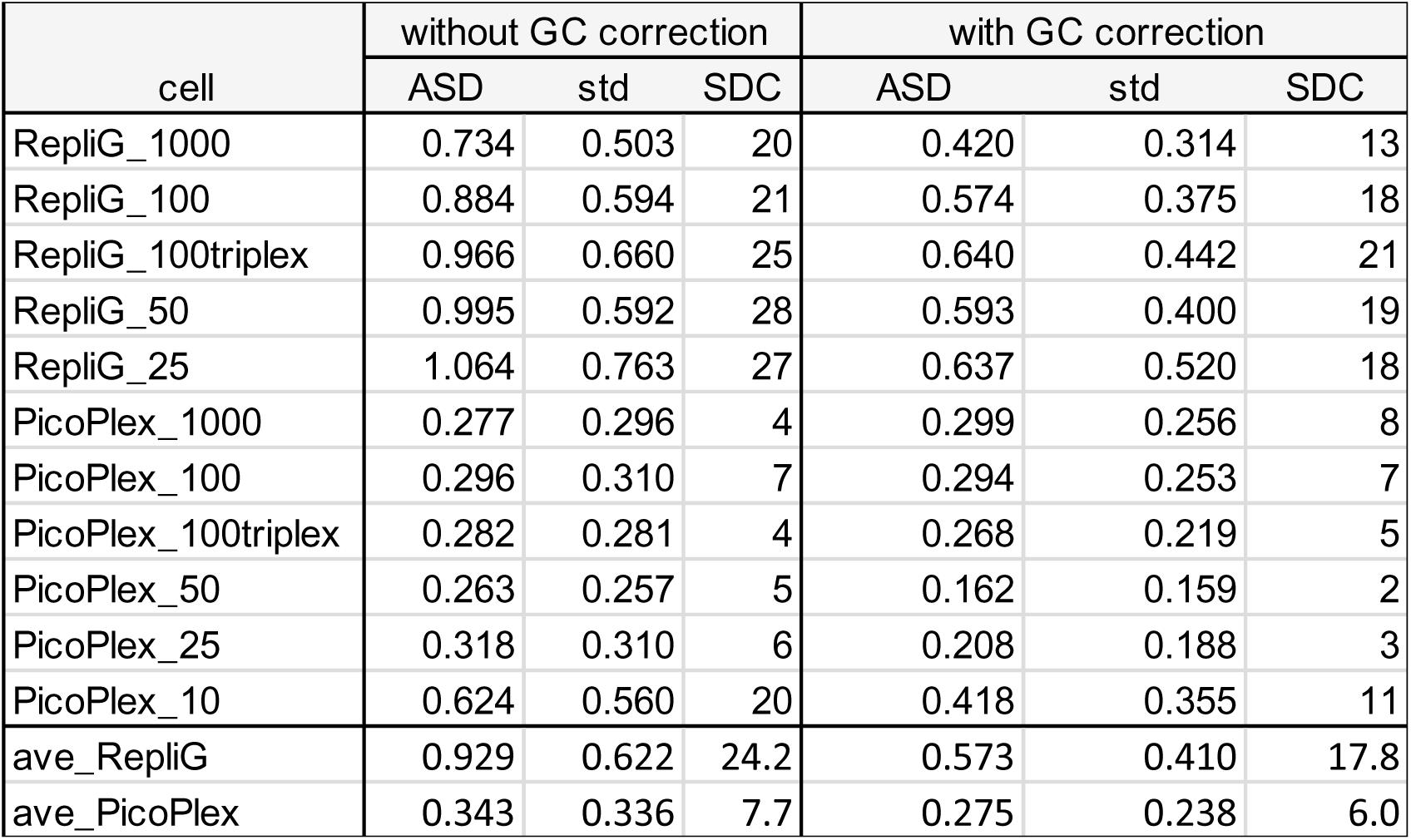
Somy estimation in RepliG and Picoplex BPK275 samples, without and with GC bias correction. ASD (average somy deviation) between a sample derived from the respective cell number (ranging from 1000 to 10) and BPK275 control and corresponding standard deviation. For somy difference count (SDC), we counted in each sample the number of chromosomes showing a somy value difference > 0.5 in comparison to BPK275 control. The average values were summarized at the bottom of the table.

In order to make an unbiased comparison on the genome coverage provided by both methods, all sequencing data sets were first subsampled to the number of reads from the sample with the lowest sequencing yield (i.e. the PicoPlex sample starting from 10 cells, containing almost 16 million reads). When comparing the fraction of the genome covered by at least one read between RepliG and PicoPlex, the former one was only slightly higher: the average ratio of those genome covering fraction for 1000, 100, 50 and 25 cells was 1.07 ± 0.05 (standard error of the ratios, Supplementary Fig. S1). When considering the fraction of the genome covered by at least ten reads, this ratio RepliG /PicoPlex increased to 1.53 ± 0.41 for these same samples. Overall a better genome coverage was thus achieved with RepliG than with PicoPlex, as visualized in the Manhattan depth plots (Supplementary data 2). This higher genome coverage is due to the fact that on average a higher percentage of reads is mapping back to the reference genome for RepliG (96.2%) compared to PicoPlex (41.1 %). However, when calculating the normalized standard deviation on the read depth – an indicator for evenness of coverage – PicoPlex shows a lower variation (Supplementary data 1, Table S1): when comparing RepliG with PicoPlex, the average ratio of those normalized standard deviations for 1000, 100, 50 and 25 cells was 1.79 ± 0.16 (standard error). This trend can be confirmed by the lower Read Count Variation of PicoPlex (see Methods). With the exception of the 1000 cells sample where both methods show a similar value, the Read Count Variation is dramatically higher for RepliG compared to PicoPlex, indicating a more even read coverage using the latter one.

The third sequence feature comparing both WGA methods was the base accuracy: this was measured by the allele frequency difference between sequence derived from a given number of cells and the control bulk DNA sample (the BPK275 control). The base accuracy was higher in RepliG samples than in PicoPlex ones, except for the sample of 10 cells (Fig. 1). The base error rates were 16.1, 9.5, 5.5, 5.1 and 4.4 fold higher in Picoplex than RepliG at cell numbers of 1000, 100, 100 triplex, 50 and 25, respectively. Only for the sample starting from 10 cells, the error rate was twice as high in RepliG compared to PicoPlex

**Figure 1.**
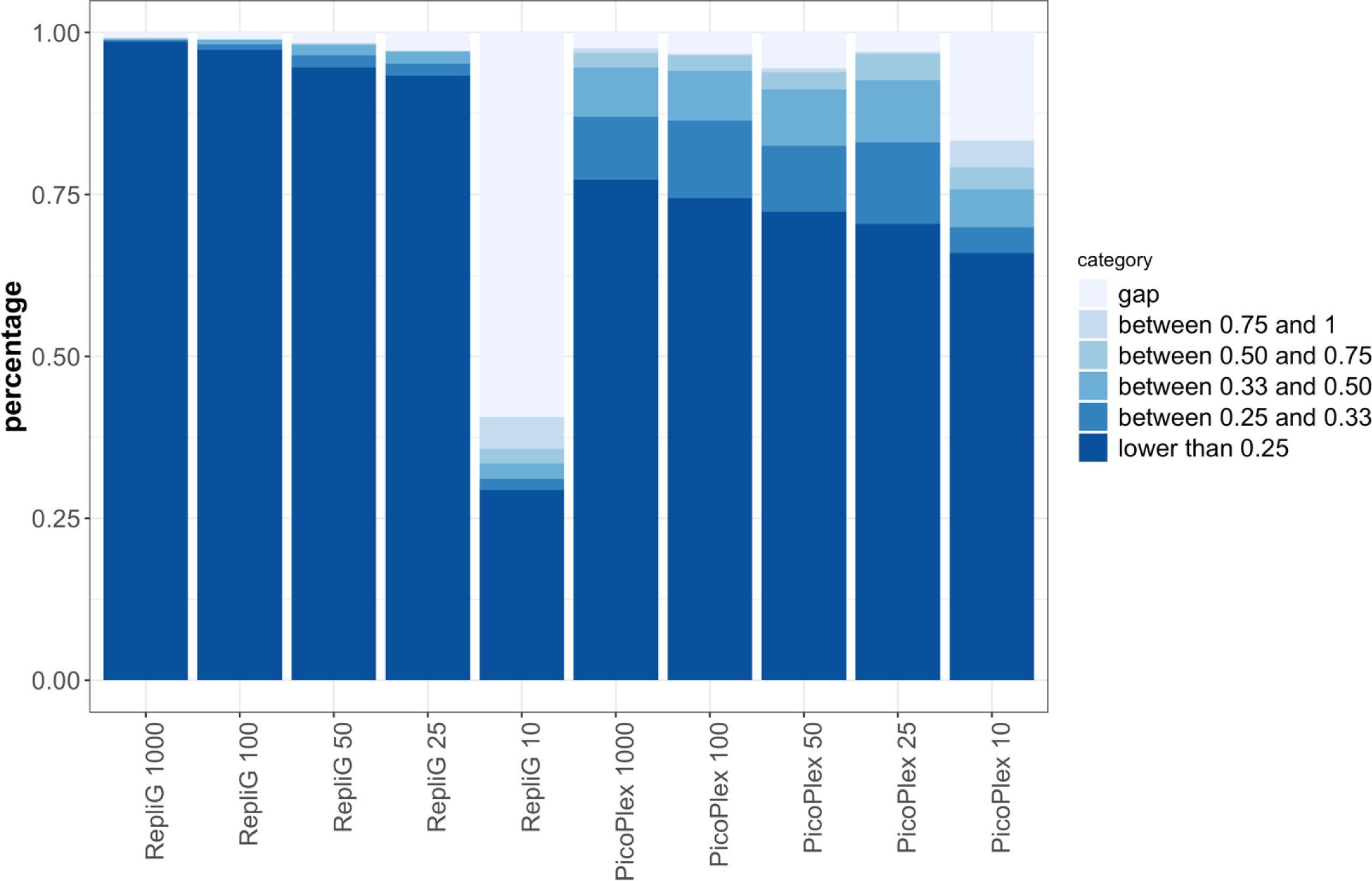
Base accuracy of the RepliG and PicoPlex BPK275 samples. Genome equivalents are shown along the x-axis. The y-axis represents the proportion of the bases with the given classifications i.e. the alternative allele frequency of the detected SNPs.

### GC bias and read depth

Both WGA methods here used were reported to be highly affected by GC content in terms of read depth distribution^14 15 16^. The GC content across the different chromosomes is expected to be uniform, but we found a strong negative correlation between the GC content and length of chromosomes in *L. donovani* BPK282 (r^2^ = 0.586, p-value: 5.47e-8, Supplementary Fig. S2 A-B) and *L. braziliensis* M2904 (r^2^ = 0.566, p-value 1.89e-7, Supplementary Fig. S2 C-D). This correlation was absent in the genomes of *Plasmodium*, *Cryptosporidium*, *Trypanosoma cruzi*, *Giardia* species deposited in EupathDB (https://eupathdb.org/eupathdb/). Lowess (locally weighted scatterplot smoothing) curves were calculated for the different samples to assess the effect of GC bias on read depth. Normalized depth only moderately depends on the GC content in the BPK275 control, resulting in a straight line with minimal slope (ANOVA on regression slope p-value < 1e-15, Tukey’s post-hoc between BPK275 control and the closest regression line (PicoPlex_1000) p-value < 1e-06). However, the effect of GC content is more pronounced in the amplified samples, with a stronger bias for RepliG treated samples than for PicoPlex treated ones (Fig.2).

**Figure 2.**
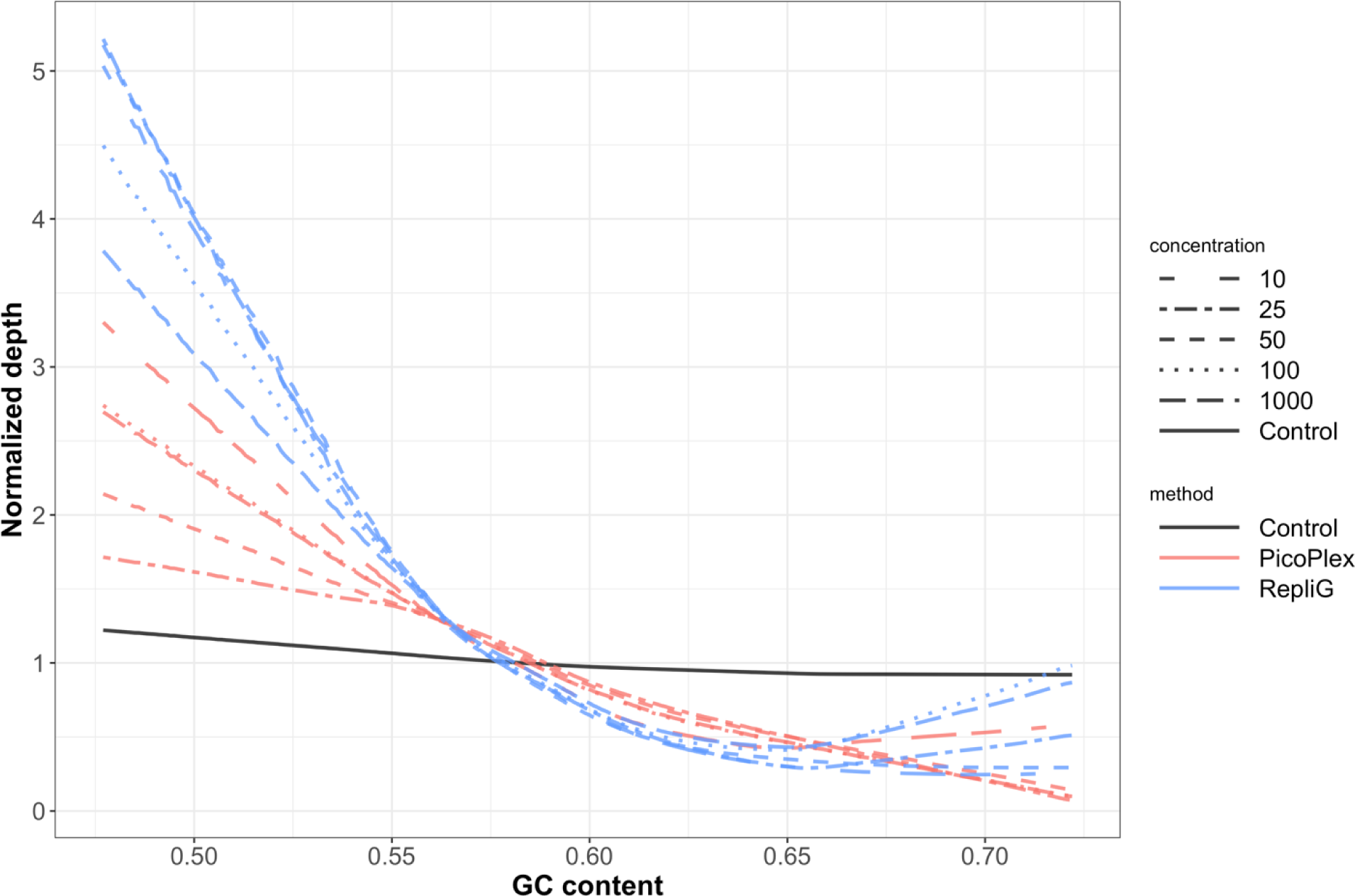
Depth dependency in relation to the GC content. Lowess curves showing the link between GC content and normalized depth for the RepliG samples (light blue), the Picoplex samples (light red) and the BPK275 control (black). The line type represents the different cell number equivalents.

### Somy determination

Boxplots were used to visualize somy values and their variation in the different samples, with and without GC correction, as described in the Methods section. The somy values of the BPK275 control had a negligible depth variability (Supplementary Fig. 3A) and the corresponding somy values corrected for GC bias were nearly identical to the values without GC correction for this control (Supplementary Fig. S4 A). For the RepliG samples, the boxplots suggest a high variability on the somy values (Supplementary Fig. 3 B). GC correction brings the somy values closer to those of the BPK275 control (Supplementary Fig. 4 B). However, somy values were still imprecise, as characterized by the average somy deviation (ASD, ranging from 0.420 to 0.637, average = 0.57) and the somy difference count (SDC, ranging from 13 to 21, average = 17.80) as shown in Table 1. In contrast, the PicoPlex samples showed smaller variation in somy values compared to RepliG (Supplementary Fig. 3 C). After GC bias correction for these samples, somy values became closer to those of the control (Supplementary Fig. 4C), which is reflected in an ASD value ranging between 0.299 and 0.418 (average = 0.28), and a SDC value ranging from 2 to 11, average = 6.00 as shown in Table 1. The graphical summary of the ASD for each sample with and without GC bias correction is given in Fig. 3 A and B, respectively. It shows (i) the lower somy deviation in PicoPlex samples (Mann-Whitney U = 0, p-value = 4.05E-03) when compared to RepliG samples and (ii) the decrease in somy deviations after the GC correction is statistically significant for RepliG samples (Mann-Whitney U = 0, p-value = 6.09E-03) but not for PicoPlex samples (Mann-Whitney U = 13, p-value = 2.36E-01). Since PicoPlex treated samples gave the most accurate somy estimates, this approach was chosen for genome amplification for single cell sequencing.

**Figure 3.**
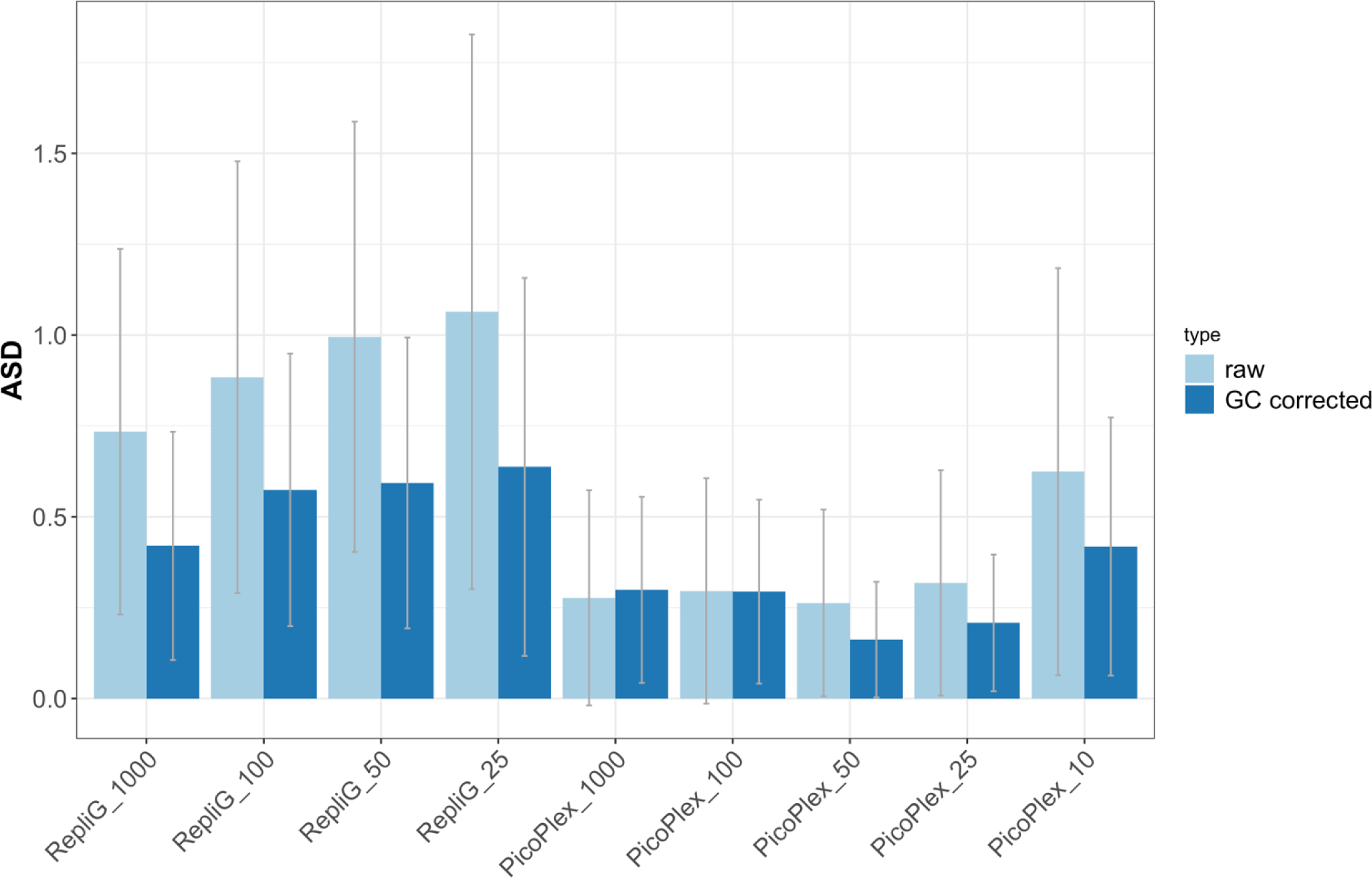
The average somy deviation (ASD) per chromosome between the BPK275 bulk control and the RepliG or the Picoplex samples, without (left bar) and with (right bar) GC bias correction. The error bars represent their standard deviation.

### Single cell genome sequencing

After comparing two genome amplification methods on DNA from different cell numbers and testing and optimizing bioinformatical methods, PicoPlex was considered more adequate than RepliG to accurately predict aneuploidy in single cells, given its more even genome coverage combined with a more accurate prediction of the real somy values. To test this in real-life single cell applications, we applied this method to a line derived from another strain, *L. braziliensis* PER094, which is characterized by a much higher degree of heterozygosity with 57,402 heterozygous SNPs than the previously used *L. donovani* BPK275 with only 43 heterozygous SNPs. This increased number of heterozygous SNPs is essential for allele frequency analyses (see below). Sorting of individual cells of the line PER094 GFP was made by FACS using the a GFP fluorescent *L. braziliensis* strain, as described in the Methods section. Two batches of cells were generated, respectively called PER094a (25 cells) and b (22 cells), each of them amplified and sequenced independently. For the batch of 25 PER094a cells, the total number of reads was 183.2 million, of which 13.3 % could be mapped to the *L. braziliensis* M2904 reference genome, resulting in average depth 2.4 ± 4.6. Similarly, for the set of 22 PER094b PicoPlex samples, 188.9 million reads were obtained of which 25.0 % of the reads could be mapped to the reference genome, resulting in average depth of 9.4 ± 10.5. All the mapping statistics of total number of reads, mapped reads and the read depth statistics for both sets of PER094 samples were given in Supplementary data 1 (Table S2). General depth trends for PER094a and b samples could be seen in Manhattan depth plots (Supplementary data 3 and 4).

**Table 2.** Estimation of somy deviation in PER094a and b single cell samples, without and with GC bias correction. ASD and SDC between a single cell sample and BPK094 control and corresponding standard deviation (see Table 1 for explanation of ASD and SDC). The single cells with even depth are shown in white and the single cells with uneven depth are shown in gray. At the bottom of the table, average of high quality (even depth, ave_pass) and low quality (uneven depth, ave_fail) are indicated. Two intermediate depth quality samples PER094a_sc7 and PER094a_sc11 (indicated in orange) were not included in the summary statistics.

| cell | without GC correction |  |  | with GC correction |  |  |
| --- | --- | --- | --- | --- | --- | --- |
|  | ASD | std | SDC | ASD | std | SDC |
| PER094a_sc19 | 0,234 | 0,147 | 2 | 0,148 | 0,093 | 0 |
| PER094a_sc20 | 0,175 | 0,130 | 1 | 0,121 | 0,104 | 0 |
| PER094a_sc8 | 0,191 | 0,129 | 2 | 0,143 | 0,111 | 0 |
| PER094a_sc9 | 0,191 | 0,145 | 1 | 0,134 | 0,116 | 1 |
| PER094a_sc6 | 0,235 | 0,159 | 2 | 0,179 | 0,122 | 1 |
| PER094a_sc24 | 0,190 | 0,155 | 2 | 0,141 | 0,124 | 1 |
| PER094a_sc5 | 0,214 | 0,149 | 3 | 0,155 | 0,125 | 0 |
| PER094a_sc18 | 0,219 | 0,157 | 2 | 0,159 | 0,127 | 0 |
| PER094a_sc13 | 0,215 | 0,142 | 2 | 0,157 | 0,128 | 0 |
| PER094a_sc14 | 0,213 | 0,158 | 2 | 0,153 | 0,135 | 1 |
| PER094a_sc15 | 0,214 | 0,161 | 3 | 0,169 | 0,142 | 2 |
| PER094a_sc23 | 0,228 | 0,151 | 2 | 0,175 | 0,142 | 2 |
| PER094a_sc21 | 0,208 | 0,156 | 1 | 0,172 | 0,143 | 2 |
| PER094a_sc7 | 0,231 | 0,178 | 3 | 0,192 | 0,169 | 3 |
| PER094a_sc11 | 0,228 | 0,182 | 3 | 0,203 | 0,191 | 2 |
| PER094a_sc2 | 0,291 | 0,293 | 5 | 0,276 | 0,252 | 4 |
| PER094a_sc10 | 0,349 | 0,402 | 7 | 0,321 | 0,424 | 6 |
| PER094a_sc17 | 2,508 | 1,622 | 33 | 1,795 | 0,833 | 32 |
| PER094a_sc16 | 2,203 | 1,412 | 32 | 2,202 | 1,413 | 32 |
| PER094a_sc12 | 0,787 | 2,203 | 8 | 0,742 | 2,240 | 6 |
| PER094a_sc25 | 2,695 | 5,016 | 14 | 2,695 | 5,095 | 11 |
| PER094a_sc3 | 1,510 | 6,154 | 10 | 1,446 | 5,872 | 9 |
| PER094a_sc1 | 1,760 | 6,680 | 8 | 1,760 | 6,680 | 8 |
| PER094a_sc22 | 3,646 | 10,454 | 11 | 3,646 | 10,454 | 11 |
| PER094a_sc4 | 4,872 | 14,477 | 11 | 4,921 | 14,666 | 11 |
| ave_pass | 0,217 | 0,162 | 2,3 | 0,167 | 0,139 | 1,2 |
| ave_fail | 2,259 | 5,380 | 14,9 | 2,170 | 5,297 | 14,0 |
|  | ASD | std | SDC | ASD | std | SDC |
| PER094b_sc17 | 0,151 | 0,133 | 1 | 0,102 | 0,105 | 1 |
| PER094b_sc7 | 0,183 | 0,152 | 2 | 0,132 | 0,120 | 1 |
| PER094b_sc1 | 0,190 | 0,148 | 2 | 0,134 | 0,121 | 1 |
| PER094b_sc15 | 0,325 | 0,209 | 6 | 0,143 | 0,139 | 2 |
| PER094b_sc13 | 0,327 | 0,209 | 7 | 0,146 | 0,135 | 1 |
| PER094b_sc12 | 0,218 | 0,144 | 1 | 0,147 | 0,122 | 1 |
| PER094b_sc19 | 0,212 | 0,164 | 1 | 0,161 | 0,140 | 1 |
| PER094b_sc16 | 0,318 | 0,219 | 7 | 0,167 | 0,152 | 1 |
| PER094b_sc20 | 0,348 | 0,237 | 10 | 0,169 | 0,172 | 2 |
| PER094b_sc9 | 0,330 | 0,216 | 5 | 0,170 | 0,183 | 2 |
| PER094b_sc5 | 0,320 | 0,222 | 6 | 0,188 | 0,153 | 2 |
| PER094b_sc3 | 0,246 | 0,160 | 2 | 0,188 | 0,139 | 2 |
| PER094b_sc4 | 0,201 | 0,179 | 2 | 0,193 | 0,153 | 3 |
| PER094b_sc2 | 0,386 | 0,203 | 10 | 0,215 | 0,173 | 3 |
| PER094b_sc22 | 0,256 | 0,382 | 5 | 0,258 | 0,418 | 6 |
| PER094b_sc8 | 0,958 | 0,659 | 23 | 0,986 | 0,606 | 26 |
| PER094b_sc10 | 1,481 | 5,473 | 8 | 1,511 | 5,626 | 9 |
| PER094b_sc18 | 1,909 | 10,174 | 3 | 1,792 | 9,571 | 2 |
| PER094b_sc6 | 7,345 | 16,187 | 10 | 7,296 | 16,173 | 9 |
| PER094b_sc14 | 14,311 | 27,530 | 16 | 14,310 | 27,530 | 16 |
| PER094b_sc11 | 18,400 | 41,837 | 15 | 19,219 | 43,741 | 14 |
| PER094b_sc21 | 20,811 | 42,345 | 12 | 21,796 | 44,363 | 10 |
| ave_pass | 0,268 | 0,185 | 4,4 | 0,161 | 0,143 | 1,6 |
| ave_fail | 8,184 | 18,073 | 11,5 | 8,396 | 18,503 | 11,5 |

Similar to the BPK275 control, the Lowess approach was used to assess the effect of GC bias on the read depth. It only showed a very limited effect for undiluted and unamplified DNA of PER094 (further called PER094 control) (Fig. 4 A). Remarkably, the PER094 control Lowess fit showed an opposite trend compared to the BPK275 control, the forming showing a positive trend with increasing GC content. However, for *L. braziliensis* single cells the GC content had a clear impact on the read depth, represented by the negative slopes as shown in Fig. 4 B and 4C for two single cells. Where the normalized depth histogram shows a normal distribution for the even depth case, the depth histogram of the uneven samples shows a peak at a normalized depth of zero, combined with a long tail distribution towards higher normalized depths, as can be seen in the dot plot and the right side depth histogram (Fig.4). This visual inspection lead to an ad-hoc cut-off of normalized standard deviation of read depth of 0.31, to distinguish between even and uneven depth samples. The Lowess curves of all the single cell samples were shown in Supplementary Fig. 5 A and B. Curves of samples with even depth (light blue) are distinct from others but they are comparable to those observed in the *L. donovani* BPK275 Picoplex samples.

**Figure 4.**
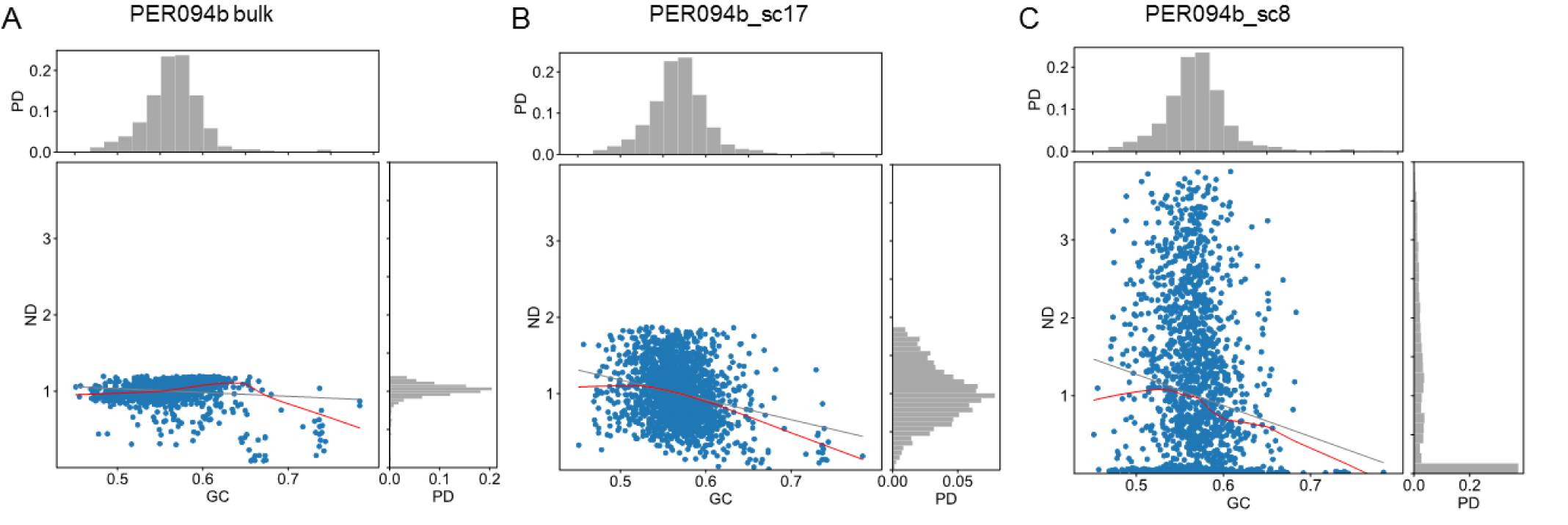
Impact of GC content on sequencing depth in the genome sequence from (A) BPK093 control, (B) and (C) a Picoplex single cell sample with even depth and uneven depth, respectively. The red lines represent the Lowess curves, the grey lines represent linear regression curves, the top and right histograms represent the histograms for the GC content (GC) and the normalized depth (NP), respectively. PD stands probability density.

In a next stage, boxplots were used to visualize the technical variability in the sequencing depth and somy estimates for all samples (Supplementary Figs 6 and 7). Variability in somy values for the PER094 control, was very limited. For the single cells, variability was higher but this was more pronounced for samples with uneven depth while somy values could be accurately determined for samples with even depth: a single dominant karyotype (called kar1) was observed in 25 single cells and it was characterized by the same trisomy of chromosomes 11 and 25 and tetrasomy of chromosome 31, while all other chromosomes were disomic. This dominant karyotype matched perfectly with the ‘average’ karyotype observed in the PER094 control. In a few other single cells with even depth, divergent karyotypes were encountered: (i) kar2, with a disomic (instead of trisomic) chromosome 25 in PER094b-sc2 and -sc9 and (ii) kar3, with monosomic chromosome 1 (instead of disomic) in PER094a-sc2 (Supplementary Figs 6, 7 and 8).

In a last step, we estimated the impact of GC bias correction on somy estimation. Among the even depth samples of PER094a and PER094b batches, GC correction could significantly lower the ASD value from 0.22 to 0.17 for PER094a cells (p-value 6.08E-05) and from 0.27 to 0.16 for PER094b cells (p-value 9.01E-05). The same trend was observed for the SDC value (2.14 and 4.4 to 1.0 and 1.6 for PER094a and PER094b cells respectively). (Table 2, Figure 5A and B). In contrast, values observed in uneven depth samples were much higher, and the effect of GC correction was negligible. The overall impact of GC correction on these samples was illustrated in Supplementary data 5 and 6.

**Fig 5.**
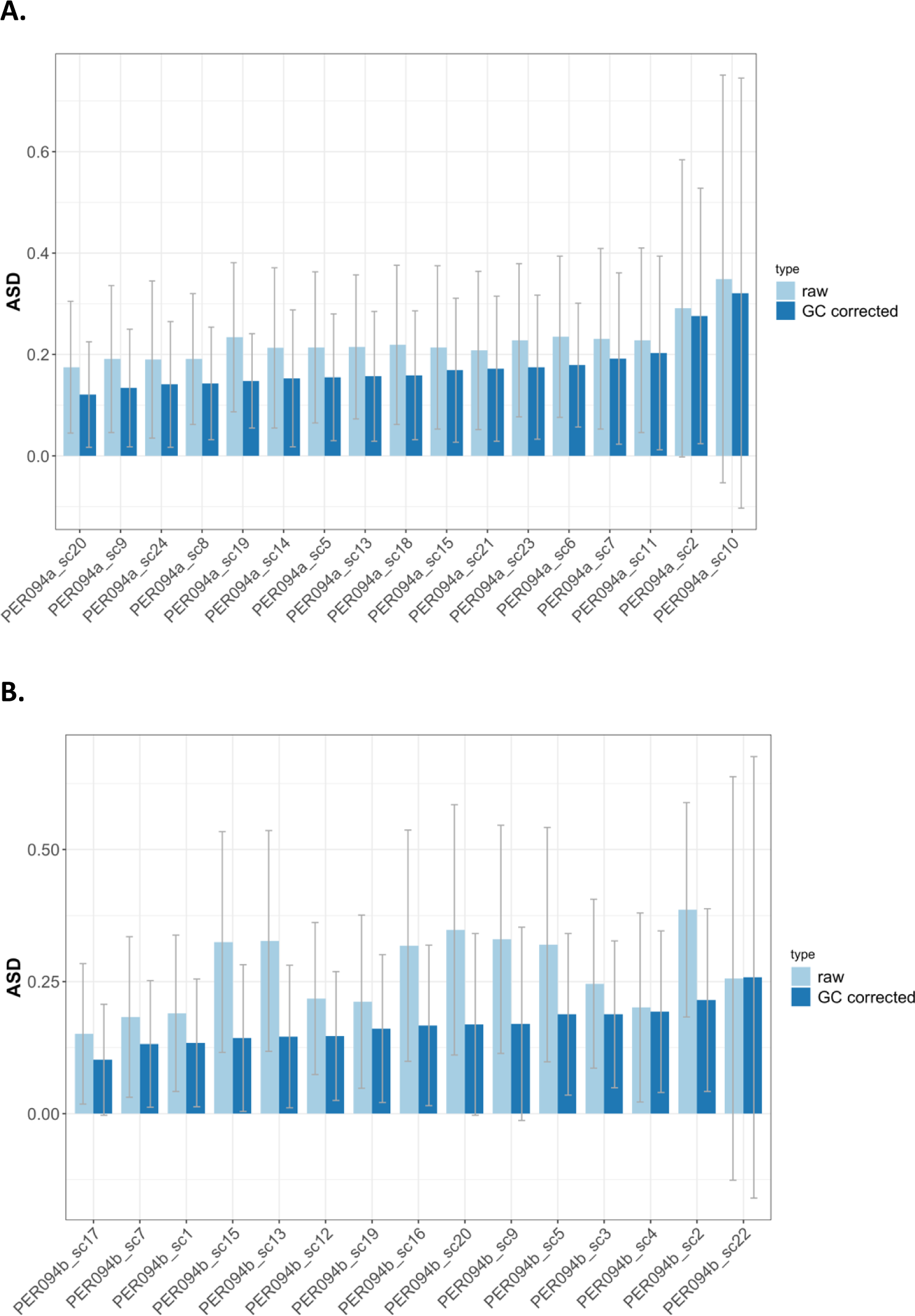
The average somy deviation (ASD) for PER094a and PER094b single cells with even depth coverage. A) ASD for PER094a single cells without (left bar) and with (right bar) GC correction. B) ASD for PER094b single cells without (left bar) and with (right bar) GC correction. The error lines represent the standard deviation. The uneven samples PER094a_sc10 and PER094b_sc22 were added as comparison at the most right side.

### Analysis of somy variation with allele frequency distribution

Somy of a given chromosome can also be predicted by the allele frequency distribution, if heterozygosity is sufficiently high^17^, which is the case for PER094. For chromosome 1, we compared the allele frequency patterns of PER094a_sc2 and PER094b_sc15, respectively shown to be monosomic and disomic by methods based on read depth (see above). We observed a typical normally distributed allele frequency of a disomic chromosome shown in PER094 control. In PER094a_sc2, however, we observed a skewed allele distribution for chromosome 1 with clear allele frequency shift toward 0 or 1 (Supplementary Fig. 9A): this was not observed for other chromosomes of that cell, hence this decrease of heterozygosity fits with monosomy of chromosome 1 in PER094a_sc2. In contrast, the chromosome 1 of PER094b_sc15 shows a disomic allele frequency pattern, leveled due to allele frequency dropout, common to single cell samples (Supplementary Fig. 9B). In the case of chromosome 25, the distribution of allele frequency is clearly bi-modal in PER094 control, which is a typical alternative allele frequency pattern of a trisomic chromosome in the bulk sequence (Supplementary Fig. 9C). Similarly, a pattern with two overlapping peaks was detected in chromosome 25 of PER094b_sc15, which is compatible with trisomy (Supplementary Fig. 9D). In contrast, the chromosome 25 of PER094b_sc9 (Supplementary Fig. 9C) showed a flat distribution similar to disomic chromosome 1 of PER094b_sc15 (Supplementary Fig. 9B).

The atypical allele frequency distribution shown above could be due to allele dropout. To verify this, we specifically measured the average pairwise base differences among a subset of PER094b PicoPlex samples (PER094b_sc20, sc15, sc2, sc13, sc9, sc5, sc16). When sites with zero depth were excluded, the average pairwise base difference at 26,959 heterozygous SNP sites was 0.303 ± 0.020 and the corresponding pairwise difference between two bulk PER094 controls was 0.020. When missing depth was treated as a base mismatch, the average pairwise base difference at 57,402 heterozygous SNP sites was 0.349 ± 0.012 and the corresponding pairwise difference between two bulk PER094 controls was 0.019. This shows that allele dropout was quite high in PicoPlex single cell samples.

## Discussion

In present paper, we report the first study – to our knowledge – of the genomes of single *Leishmania* cells. Using *L. donovani* as model, we showed in a first step that the choice of WGA method should be determined by the planned downstream genetic analysis and that Picoplex and RepliG are more suitable for aneuploidy and SNP calling, respectively, as expected from previous work^10^. Given our interest in aneuploidy^5, 6, 18^ and mosaicism^19^, we evaluated and developed bioinformatic methods to assess the somy for each chromosome when only limited sequencing data is available, thereby taking into account and correcting for the GC-bias. In a second step, we used the PicoPlex approach to sequence the genome of FACS-sorted single cells and determined their aneuploidy by the computational pipeline optimized in the first step. For this experiment, we used *L. braziliensis* as model, as its higher heterozygosity should permit to study allele frequency distribution, which is an complementary inferential method for somy estimation often applied in bulk sequencing^17^. We successfully called somy of all 35 chromosomes in 28 out of the 47 single cells, detected aneuploidy mosaicism by read-depth based methods and subsequently validated monosomy and trisomy of some chromosomes based on their signatory somy specific allele frequency distribution.

Bioinformatic pipelines were optimized to handle high technical read depth variation observed throughout the genome because of the required amplification process of WGA. We have developed and validated on low DNA amounts depth normalization methods. We tested a various range of window sizes and percentiles to quantify sequencing depth, greatly enhancing the sensitivity and specificity for somy detection. It also appeared critical to select the lower and upper depth bin cut offs for somy estimation; accordingly, these parameters must be optimized for each data set separately. In general we found that the combination of higher read depth and even read distribution always led to improved somy estimation based on depth and read allele frequency. Samples with even but low depth could still lead to accurate somy estimation. However, samples with higher intra-chromosomal read depth variation, regardless of its read depth size, always led to very poor somy accuracy.

Previous *Leishmania* studies based on bulk sequencing have shown some correlation between read coverage depth and chromosome length^6, 17, 20^. In a previous study where we used SureSelect, a genome capture method, we observed the tendency that the shorter chromosomes had lower depth which resulted in more non-integer somy values compared to longer chromosomes in the sequencing results^21^. This bias was corrected using sequencing depth normalization using chromosomes with similar lengths. The observation of non-integer somy values in smaller chromosomes could be explained by a higher mosaicism among this size-class of chromosomes^17^. However, present study provides an alternative answer to these observations, as we found in reference genomes of both *L. donovani* and *L. braziliensis* a negative correlation between GC content and chromosome length. Interestingly, this appeared to be a *Leishmania*-specific feature as this GC-bias was not found in other tested genomes. The detection of this GC-bias has an impact for single-cell genomics of *Leishmania*, because WGA methods are reported to be particularly sensitive to GC content^14–16^. In our analysis, the slopes of the Lowess curves, used to represent the link between normalized depth and GC content were bigger in *L. donovani* RepliG samples than in PicoPlex ones (Fig. 2). Accordingly, GC correction appears to be important for single cell genomics of *Leishmania*.

In this study we sorted cells using FACS and we amplified and sequenced separately single genomes. Major disadvantages, previously reported to be associated with a FACS-based approach, are the high risk of contamination with foreign DNA during collection process and the high cost of WGA of each sample. First, the size of the microbial genome is much smaller than mammalian genomes and even small fragments of this DNA will efficiently compete with the microbial DNA during WGA. This was clearly illustrated for bacteria^22^ and could represent an issue for *Leishmania* genome which is 100x smaller than human genome. We used here a FACS instrument, which was cleaned and sterilized, but not with a DNA removing buffer. Second, the high cost of WGA in FACS-based approaches is related to the volume of sample collection (5 µl here) and reaction (75 µl here) and this may also have an effect on the efficiency of the amplification of minute amounts of DNA in the sample ^22^. In our experiments, we tested the amplified *Leishmania* genomes for presence of human DNA by qPCR of only one human gene, RPL30 (see Supplementary methods). The presence of this gene was detected only in few samples, and these were excluded from further studies. However, traces of contamination with human DNA were still detected by WGS, but this was not found to be a critical factor for the somy accuracy. Indeed, a large proportion of unmapped reads did not necessarily lead to the lower somy calling accuracy. Reciprocally, a large proportion of mapped reads did not necessarily lead to higher somy calling accuracy, the latter mainly because this type of samples often showed an uneven genome coverage. Using automated, microfluidics- or droplet-based single cell sorting and preparation platforms would greatly reduce this risk and increase reproducibility of single cell whole genome analysis. Among the additional advantages of these methods, we mention: 1) the volume of the reaction (a few nanoliters), decreasing risk of contamination and cost and increasing reaction’s efficacy; 2) the use of chips, which decreases the number of operations and the risk of contamination; 3) pre-staining or constructing parasites harboring a fluorescent marker is not needed, thus most types of unprocessed cells can be analyzed without lengthy preparation; 4) the high-throughput character of analyses and 5) the possibility -with some platforms- to check the number of cells present in each sample and sequence only the ones containing only one cell, hereby also reducing costs and ensuring that only individual cells are analyzed.

Despite the low number of *L. braziliensis* cells analysed here (28 with even depth), mosaicism could be detected, with 3 different karyotypes, all aneuploid: one dominant karyotype (kar1) in 25 cells and two others (kar2 and kar3) each one encountered in 2 and 1 cells respectively. Interestingly, kar2 and kar3 only differed slightly from kar1, by the somy of 1 and 2 chromosomes respectively. Chromosome 31, shown to be tetrasomic in bulk sequences of all *Leishmania* species studied so far^5, 17, 20^ was tetrasomic too in all 28 cells analyzed here. Probably because of allele drop-out, rather high in Picoplex samples, allele frequency distribution curves of individual chromosomes (expected to be monomodal, bi-modal and tri-modal for disomic, trisomic and tetrasomic chromosomes) were atypical in single cells. However, we could detect the allele frequency signatures of monosomic chromosome 1 and trisomic chromosome 25 and thus validate the read-depth based somy calling of these chromosomes. As such, our single cell sequencing data confirm the hypothesis of mosaic aneuploidy which was derived based on FISH data.

This study paves the way for further single cell genomics studies in *Leishmania*. The FACS-based approach described here is of interest for in-depth analysis of genomes in a small number of cells (for instance a plate of 96 cells), while different WGA methods should be used depending on the planned downstream genetic analysis (SNP, indel, CNV or aneuploidy). High-throughput analyses of single cells are needed to investigate the extent and dynamics of aneuploidy mosaicism in *Leishmania*, in both stable and variable experimental conditions. Therefore, microfluidics- and droplet-based platforms represent a promising alternative and several options exist (see recent review^23^).

## Materials and Methods

### Preparation of samples for comparison of WGA methods

The cloned line *L. donovani* MHOM/NP/03/BPK275/0-Cl18^20^ was grown on HOMEM and harvested 20 passages after cloning. Parasites were washed and resuspended in PBS, to have 1000, 100, 50, 25 and 10 promastigotes in 2 µl of PBS. For Picoplex (New England Biolabs), 2 µl of parasites were put in PCR tubes and flash-frozen in liquid nitrogen. For RepliG (Qiagen), 2 µl of parasites + 2 µl of PBS were put in PCR tubes and flash frozen in liquid nitrogen. Picoplex and RepliG samples were then further processed according to manufacturer’s instructions. The average size of the DNA fragments after amplification differed between RepliG and PicoPlex, being 2-100 kb and 0.1-1 kb, respectively. DNA of each sample was purified and concentrated with Genomic DNA Clean and Concentrator-25 (Zymo Research) according to the manufacturer’s instructions and samples were frozen at −20°C. For both RepliG and PicoPlex, 100triplex samples were prepared for replicability control: for this, 100 cells were amplified in triplicate and then pooled for sequencing. DNA from the same *L. donovani* line and the same batch was extracted with the QIAamp DNA Mini Kit (Qiagen) according to manufacturer’s instructions and used as bulk control (further called BPK275 control). Samples were sent to WTSI for whole genome sequencing: (i) in 40µl, with a DNA concentration ranging between 63.3 and 91.3 ng/µl for RepliG samples, (ii) in 34 µl, with a DNA concentration ranging between 9.64 and 40.4 ng/µl for PicoPlex samples and (iii) in 40 µl, with a concentration of 50.4 ng/µl for BPK275 control.

### Preparation of samples for single cell analysis

A transgenic line of *L. braziliensis* strain MHOM/PE/02/PER094 with constitutive expression of the GFP reporter integrated within the ribosomal locus was generated as previously reported elsewhere^24^. After transfection and two weeks of selection a clonal/isogenic line was derived by the micro drop method. The resulting GFP fluorescent line PER094-GFP-Cl2 (further called PER094-GFP) was used to sort single cells within 96 well plates with the BD FACSAria II with a 85 μM nozzle and 45 PSI. Briefly, 2 ml of parasites were washed with 10 mL of PBS. Subsequently, the pellet was resuspended in another 10 mL of PBS and gently passed through a 5 μM filter (pipetting and gravity). The recovered cells were concentrated by centrifugation (1500 g, 5 min) and brought to a new suspension of 20 x 10^6^ parasites/mL in medium M199 + 100 units/mL of penicillin and 100 μg/mL of streptomycin. For sorting the single cells, gates were selected by using the side and forward scatter plots. A non-GFP wild type was included to stablish the autofluorescence values and the gate corresponding to GFP positive cells. The single cells were sorted in a 96 well plate (containing 5 µL of lysis buffer) and immediately place on ice until the next step. In parallel, DNA from the same PER094-GFP line and the same batch as the one used for single cells was extracted with the QIAamp DNA Mini Kit (Qiagen) according to manufacturer’s instructions and used as bulk control (further called PER094 control).

Two sorting experiments were made and generated 2 batches of 39 and 40 single cells. A quality control was introduced before further selecting cells for WGA and WGS. DNA contained in each well was amplified by 4 qPCR assays targeting (i) kDNA, 18s rDNA and G6PD to assess for the presence of good quality *Leishmania* DNA and (ii) RPL30 to assess for the presence of possible human DNA contamination (supplementary methods). After this process, the 2 batches resulted in 25 and 22 cells, further called PER094a and PER094b, respectively.

The performance of the cell sorter was also evaluated by sorting fluorescent beads (10 μM, Beckman Coulter) within 384 plates (optically clear flat bottom, Perkin Elmer). The presence of only one bead was corroborated by visualization of the whole well with a confocal microscope (Zeiss LSM 700). The percentage of wells with only one bead was (78.8) while the percentage of wells without or with more than one bead were 20.8 and 0.4 % respectively.

### Whole genome sequencing

For Illumina sequencing genomic DNA was sheared into 400–600 base pair (bp) fragments by focused ultrasonication (Covaris Adaptive Focused Acoustics technology, AFA Inc., Woburn, USA). Amplification-free Illumina libraries were prepared^25^ and for PER094 samples, 151 bp paired-end reads were generated on an Illumina HiSeq X, while for BPK275 samples 100bp paired-end reads on HiSeq v4 following the manufacturer’s standard sequencing protocols^26^, but with 10% PhiX DNA added to each library.

### DNA read mapping, SNP calling

Paired-end reads from the 6 RepliG and the 6 Picoplex samples of *L. donovani* and from BPK275 control were mapped to the improved reference *L. donovani* genome LdBPKv2^6^ (available at ftp://ftp.sanger.ac.uk/pub/project/pathogens/Leishmania/donovani/LdBPKPAC2016beta/) using Smalt v7.4 (http://www.sanger.ac.uk/science/tools/smalt-0). Similarly for *L. braziliensis*, paired-end reads from 22 PER094b and 25 PER094a FACS-sorted single cells and the PER094 control were mapped to the *L. braziliensis* M2904 reference genome, which was recently improved based on PacBio SMRT sequencing. The parameters used in Smalt were described in the previous studies^6, 21^. For calculation of the genome coverage for the different samples, samtools was used to subsample all data to the amount of reads obtained for the sample with the lowest yield. To assess the evenness of genome coverage, the variation on the genome coverage within 5kb windows was calculated using the normalized standard deviation (also called coefficient of variation), i.e. the standard deviation of the sequencing depth within 5kb windows divided by the average sequencing depth calculated over all 5kb windows. The read count variation was calculated as specified using 5kb windows^27^.

For SNP calling for the evaluation of the accuracy of sequences derived from amplified and bulk DNA, we used the population SNP calling mode of UnifiedGenotyper in Genome Analysis Toolkit v3.4, with the SNP cut off (GATK QUAL score) (i) 1500 for BPK275 samples and (ii) 2000 for PER094 samples (GATK: https://software.broadinstitute.org/gatk/)^28^. The SNP cut off for *L. braziliensis* was higher since more samples were used to call SNPs. The main goal of the SNP analysis in this study was to evaluate base accuracy and the base recovery rate, i.e. how many bases would be covered by at least one read. Therefore, no lower nor higher depth cut off was imposed in the population SNP calling. In *L. donovani* chr22, the base positions between position 736224 and the end of that chromosome were excluded from the SNP calling since many false positive heterozygous SNPS were found in this region.

### GC content and GC bias correction

With WGA methods like RepliG and PicoPlex, normalized depth is reported to be affected by GC content bias^14–16^. Previous Leishmania studies based on bulk cell populations have suggested a correlation between read coverage depth and chromosome length ^2029^. However, the mechanism of depth bias was not fully understood. Neighboring chromosome normalization ^29^ was used to assess this effect. However, this will not be accurate for samples in which the variability of chromosome copy number is irregular and skewed. First the GC content was measured in 5 kb windows across each chromosome. To evaluate the impact of GC content bias on the current data sets, we selected the long disomic chromosomes 28, 29, 30, 32 and 34 of *L. donovani* and *L. braziliensis*. These disomic chromosomes were used to avoid the depth difference due to aneuploidy. For each sample, we fitted a Lowess (Locally Weighted Scatterplot Smoothing) curve in terms of the GC content and normalized depth, using a Python package statsmodels v0.9.0 (https://github.com/statsmodels/statsmodels). Depths greater than the 95 percentile were marked as outliers and removed.

In the next step this Lowess curve is used to correct the somy value for each chromosome separately. Based on the average GC content of a chromosome, its somy value is divided by the value of the Lowess curve for that specific GC percentage using a lookup hash table approach. In general, the Lowess curves properly represented the relationship between the GC content and depth within a GC content range between 55% and 65% for most of the high-quality samples. As the average GC content of each chromosome was between 56.2% and 61.5% for both *L. braziliensis* and *L. donovani* genomes, accurate GC correction could be guaranteed. Finally, somy values were renormalized using a median somy value after the initial GC correction to have the median somy value over all chromosomes close to 2. The method used here was analogous to the GC bias correction methods described in the previous studies^30, 31^. Somy values with and without the GC correction were visualized with Gnuplot^32^.

### Somy estimation

WGA methods are known to produce highly variable depth coverage^14, 16^. To mitigate skewed, uneven read distribution^6^, the chromosome median depth was calculated as a trimmed median of mean depths for 5000bp bins, where bins with a depth less than the 10^th^ percentile or greater than 90^th^ percentile were removed. Somy values are visualized using boxplots as implemented using Matplotlib^33^. To assess depth across all the chromosomes, Manhattan plots across all the chromosomes were created based on median depth of 5000bp windows. The upper limit of a Manhattan plot was set to be twice the value of a 98 percentile to focus on the informative range.

### Somy range

The range of monosomy, disomy, trisomy, tetrasomy, and pentasomy was defined to be the full cell-normalized chromosome depth or S-value: S < 1.5, 1.5 ≤ S < 2.5, 2.5 ≤ S < 3.5, 3.5 ≤ S < 4.5, and 4.5 ≤ S < 5.5, respectively^6^. However, single cell sequencing depth can be highly variable among cells unlike in bulk sequencing. To overcome this technical depth variability among individual cells and to characterize somy variability clearly, we defined the Average Somy Deviation (ASD): this is defined as the average difference between the calculated somy value and the true (integer) somy value, with the latter one calculated based on the bulk control sample. Similarly, the somy difference count (SDC) is defined as the number of chromosomes where the absolute difference predicted somy value and the true somy value is greater than 0.5 .

### Visualization of pairwise allele frequency difference

Alternative base allele frequencies (allele frequencies) of variable sites were extracted from the GATK SNP vcf files. To quantify base recovery rate, a gap was considered to be homozygous mismatch of allele difference of one, instead of discarding missing sites or imputing the bases. For *L. braziliensis* single cells, we were particularly interested in identifying allele frequency shifts due amplification artifacts. Therefore we first identified high quality heterozygous sites in the PER094b and PER094a bulk samples, and only assessed allele frequency differences. We did not take this approach for the *L. donovani* samples since there are not enough heterozygous SNPs in BPK275. An allele frequency distribution of two samples was visualized with an allele frequency dot plot where the x- and y-coordinates represented their alternative allele frequencies, and their corresponding histogram was also given along each axis (Supplementary Fig. 9).

## Acknowledgements

This study received financial support from the Flemish Ministry of Science and Innovation (SOFI Grants SINGLE and MADLEI), the Flemish Fund for Scientific Research (FWO, post-doctoral grant to MJ). JAC, MB and MS were supported by Wellcome via the core support of the Wellcome Sanger Institute (grant 206194).

## Author contributions

All authors have approved the submitted version of this manuscript and have agreed both to be personally accountable for their own contributions and to ensure that questions related to the accuracy or integrity of any part of the work are appropriately investigated and resolved.

This work was conceived and designed by HI, MB, JAC, MV, JCD and MAD. Data were acquired and analyzed by HI, PM, MJ, MS, IM, JAC and MAD. Data interpretation was made by HI, PM, MJ, JAC, JCD and MAD. Paper was drafted by HI, PM, JCD and MAD and substantively revised by MJ, MS, MV, MB and JAC.

## Data availability

Sequencing data are available in ENA (European Nucleotide Archive) study PRJEB8793

## Supplementary information

**Supplementary Figure 1.**
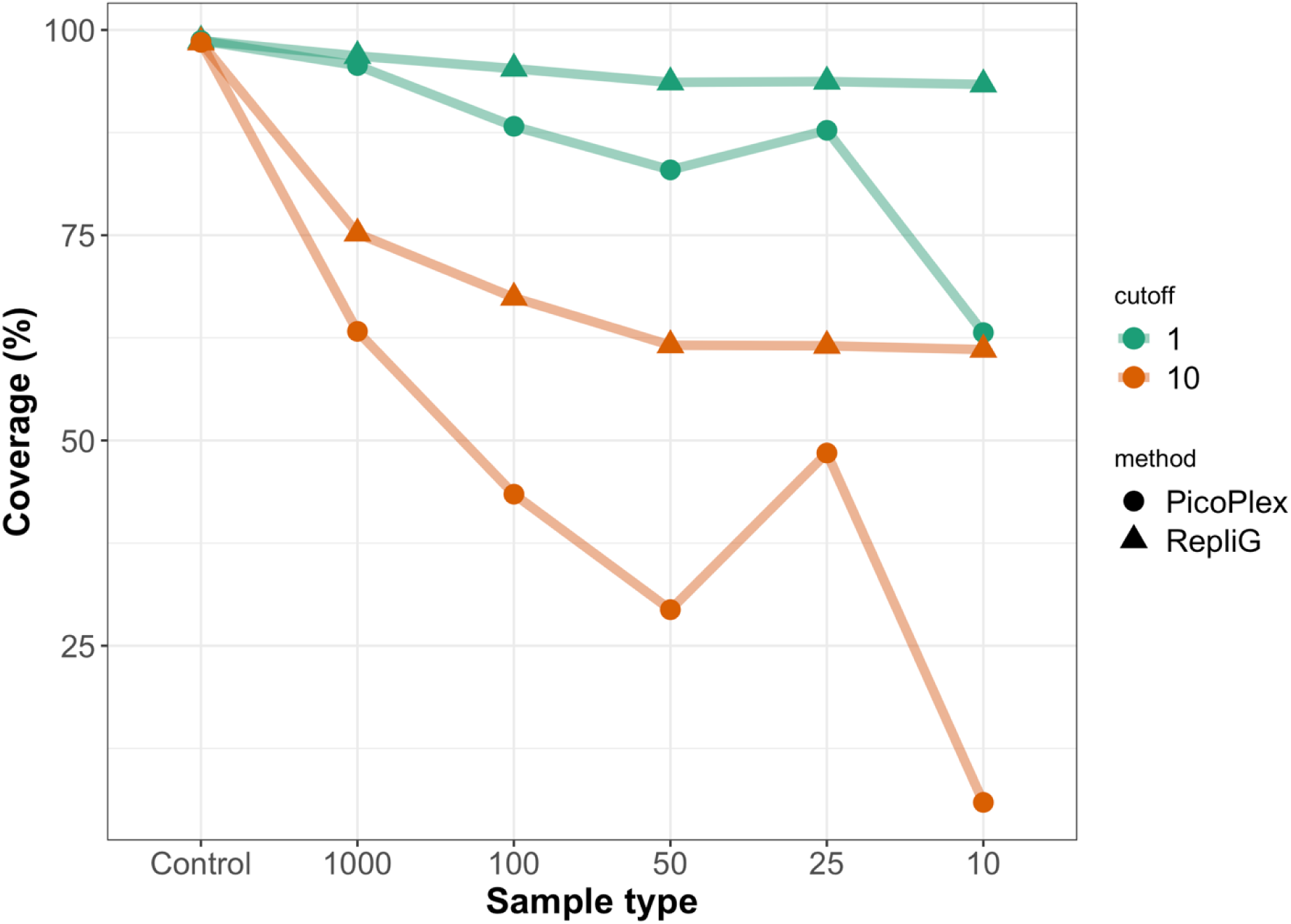
For DNA extracted from different amounts of *L. donovani* BPK275 cells (1000, 100, 50, 25 or 10 cells), and amplified using RepliG or PicoPlex, the fraction of the genome covered by at least 1 read (green line) or 10 reads (red line) is plotted. The control sample is obtained from undiluted and unamplified DNA.

**Supplementary Figure 2.**
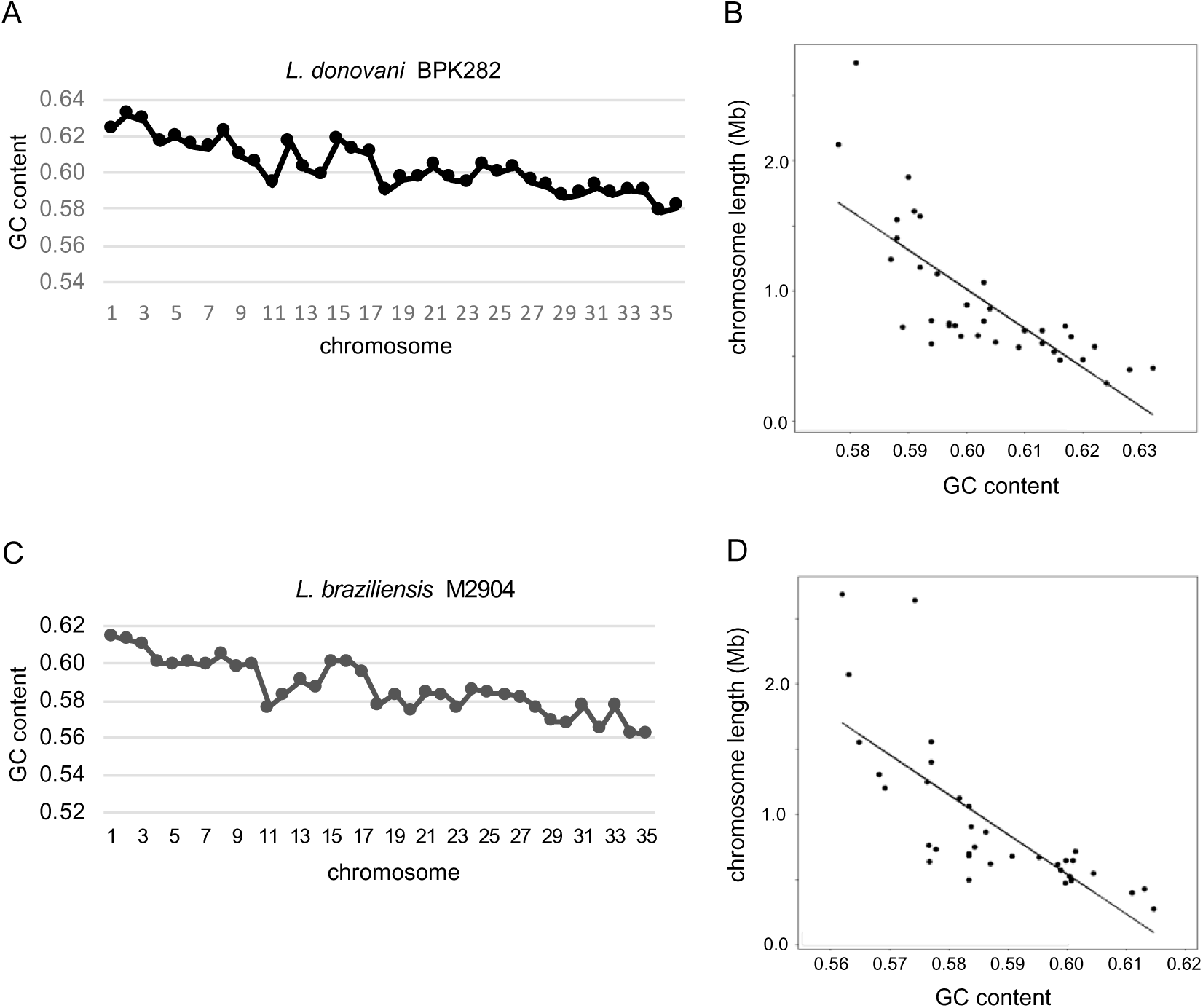
Chromosome length and GC content in reference genomes of *L. donovani (Ld)* BPK282 (A) and *L. braziliensis* (*Lb*) M2904 (C). The GC content was plotted for each chromosome. Correlations: (B) for *Ld*, r^2^ = 0.586, p-value 5.47e-8, (D) for *Lb*, r^2^ = 0.566, p-value 1.89e-7.

**Supplementary Figure S3.**
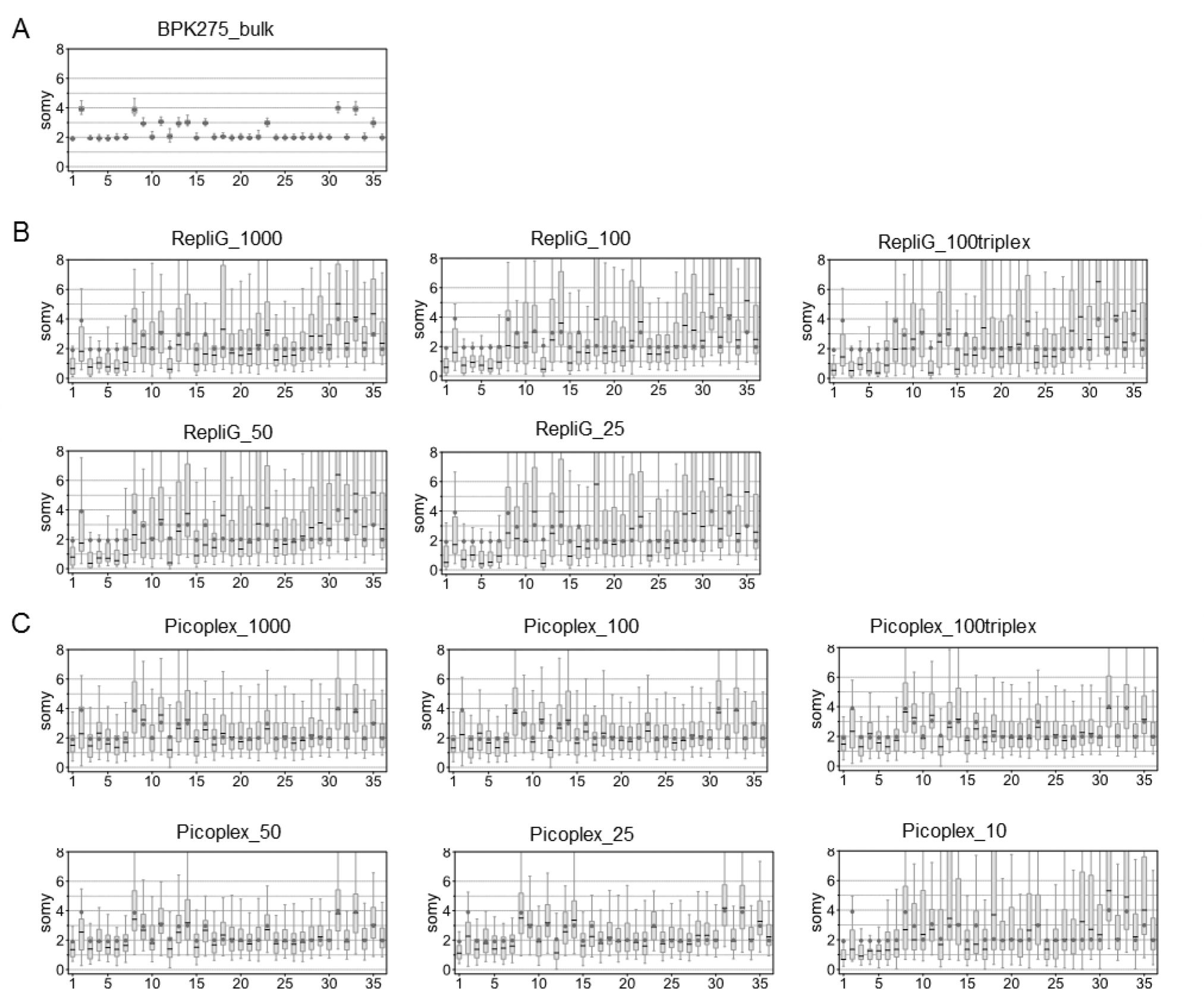
Box plots for visualization of somy estimations, where a somy estimate is calculated for each 5kb window in (A) BPK275 control, (B) RepliG samples derived from different cell numbers and (C) PicoPlex samples derived from different cell numbers. The true somy value – derived based on the BPK275 control – is given in a grey filled circle. The median somy of a chromosome in a given sample is shown as thick horizontal line.

**Supplementary Figure S4.**
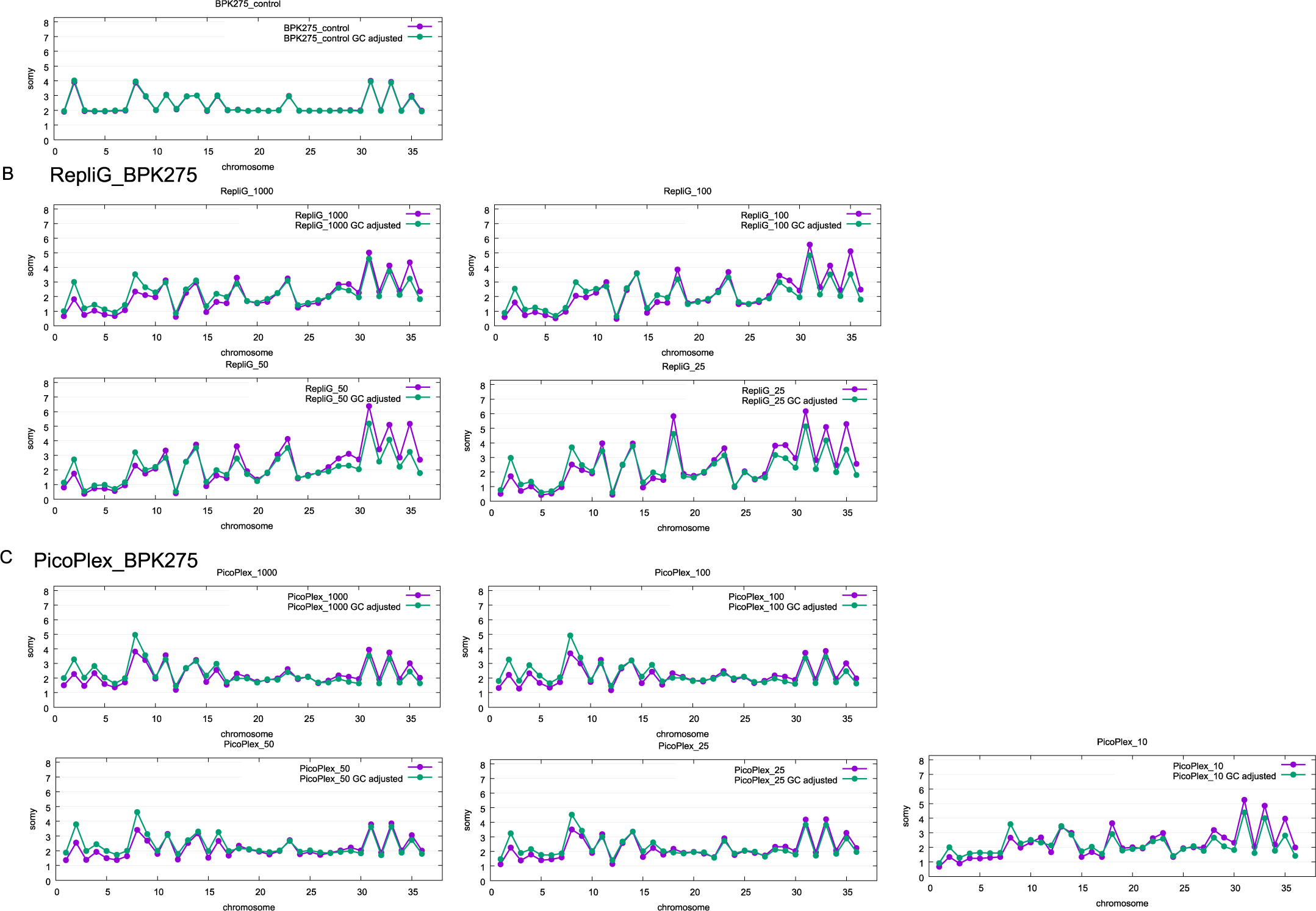
Somy visualization without (purple line) and with (green line) GC bias correction in (A) BPK275 control, (B) RepliG and (C) Picoplex samples. The x-axis and the y-axis represent chromosome and somy value, respectively.

**Supplementary Figure S5.**
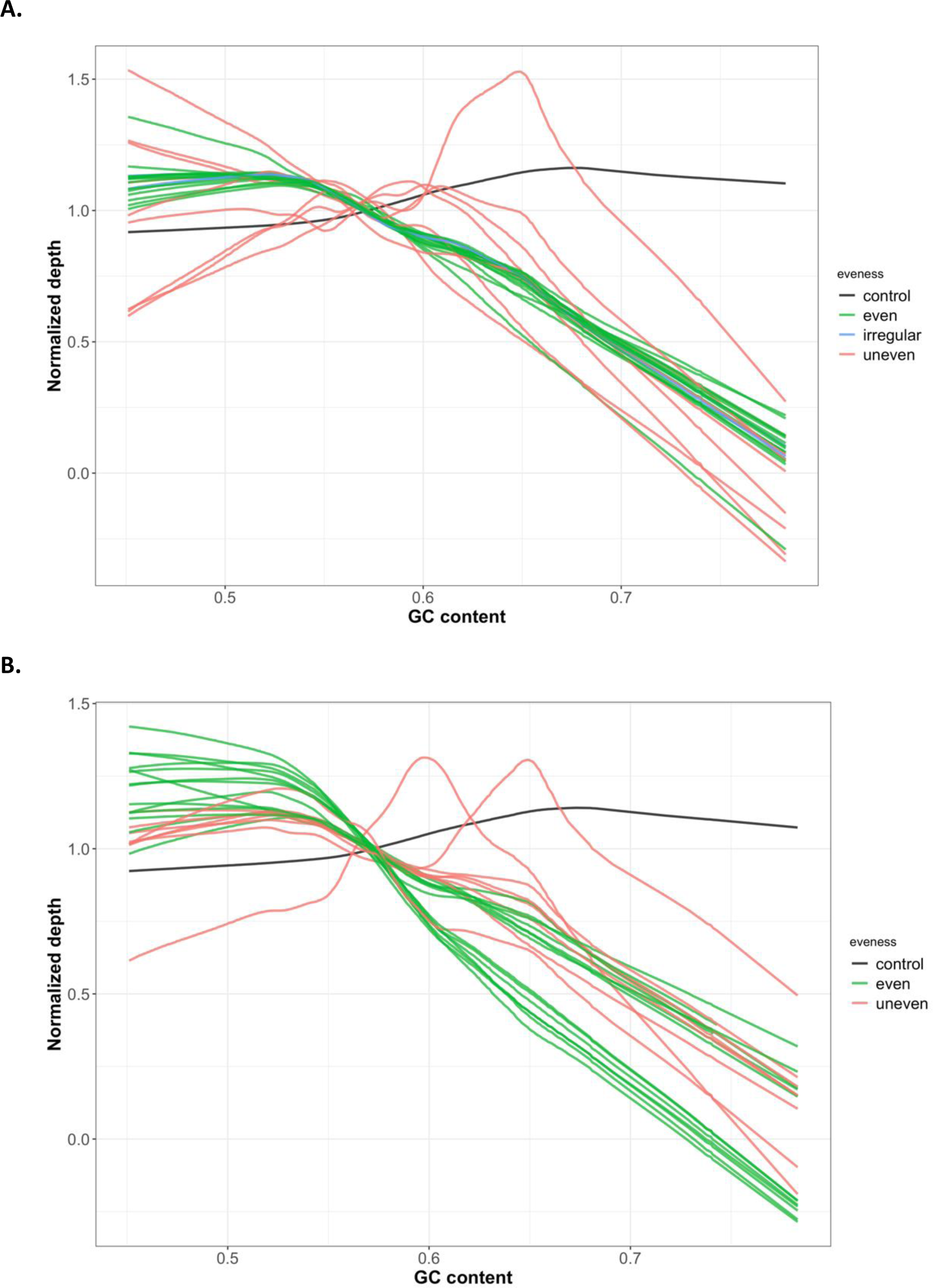
Lowess curves showing the impact of GC content on normalized depth in (A) PER094a samples and (B) PER094b samples. The bulk controls are shown in black. The even, mixed and uneven depth samples were shown in light blue, light green and light brown, respectively.

**Supplementary Figure 6.**
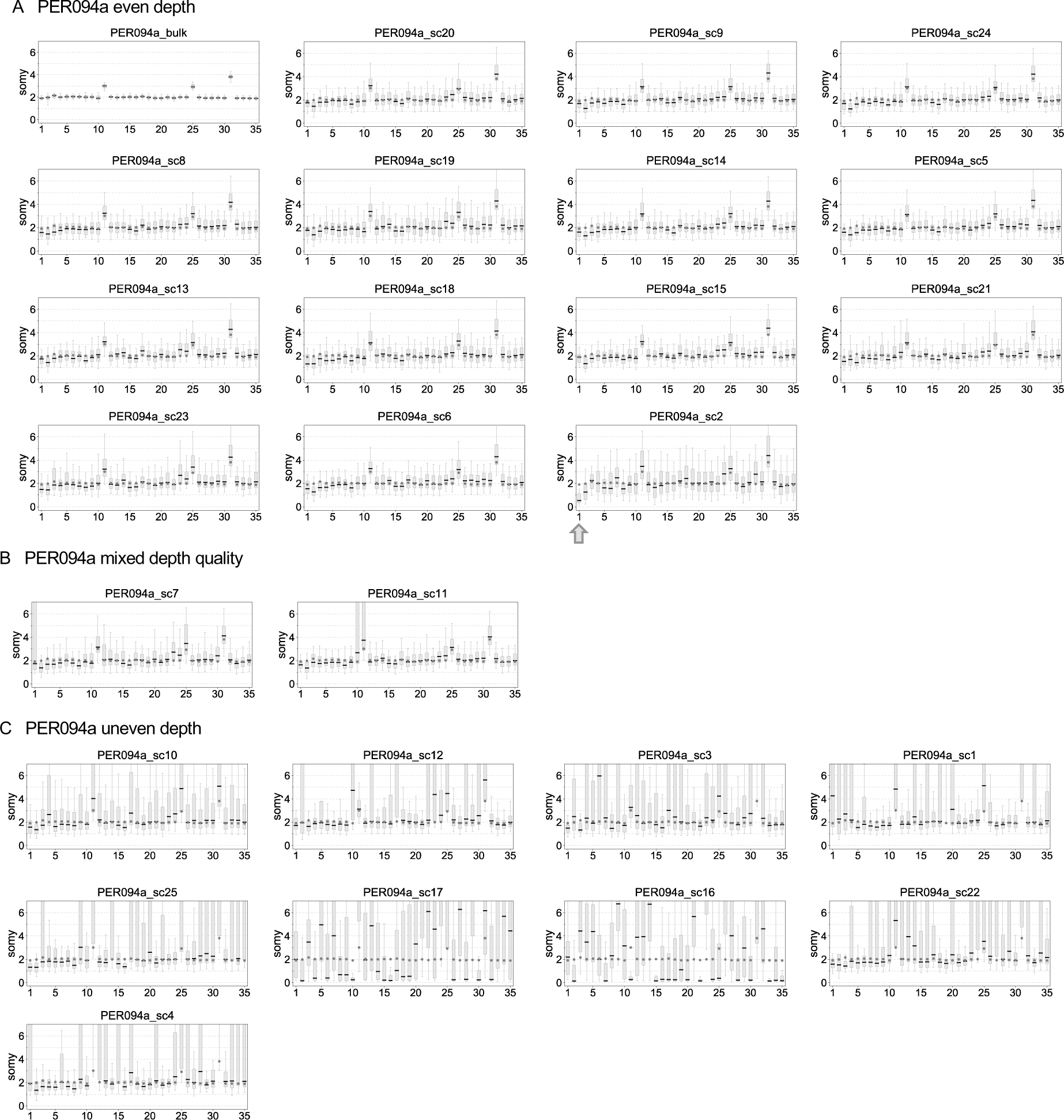
Boxplots of PER094a single cell samples: A) The first boxplot was for the PER094 control. There were 14 even depth single cells that had a short box and short error whiskers, allowing accurate somy estimation. Chromosome 1 is disomic in most samples, except PER094a_sc2, where it is monosomic, marked by the arrow. B) Mixed depth: PER093A_sc7 and PER093A_sc11 had 1 and 2 chromosomes respectively, with high variable depth. C) The uneven depth of 9 samples led to an erratic somy estimation that could not be corrected.

**Supplementary Figure S7.**
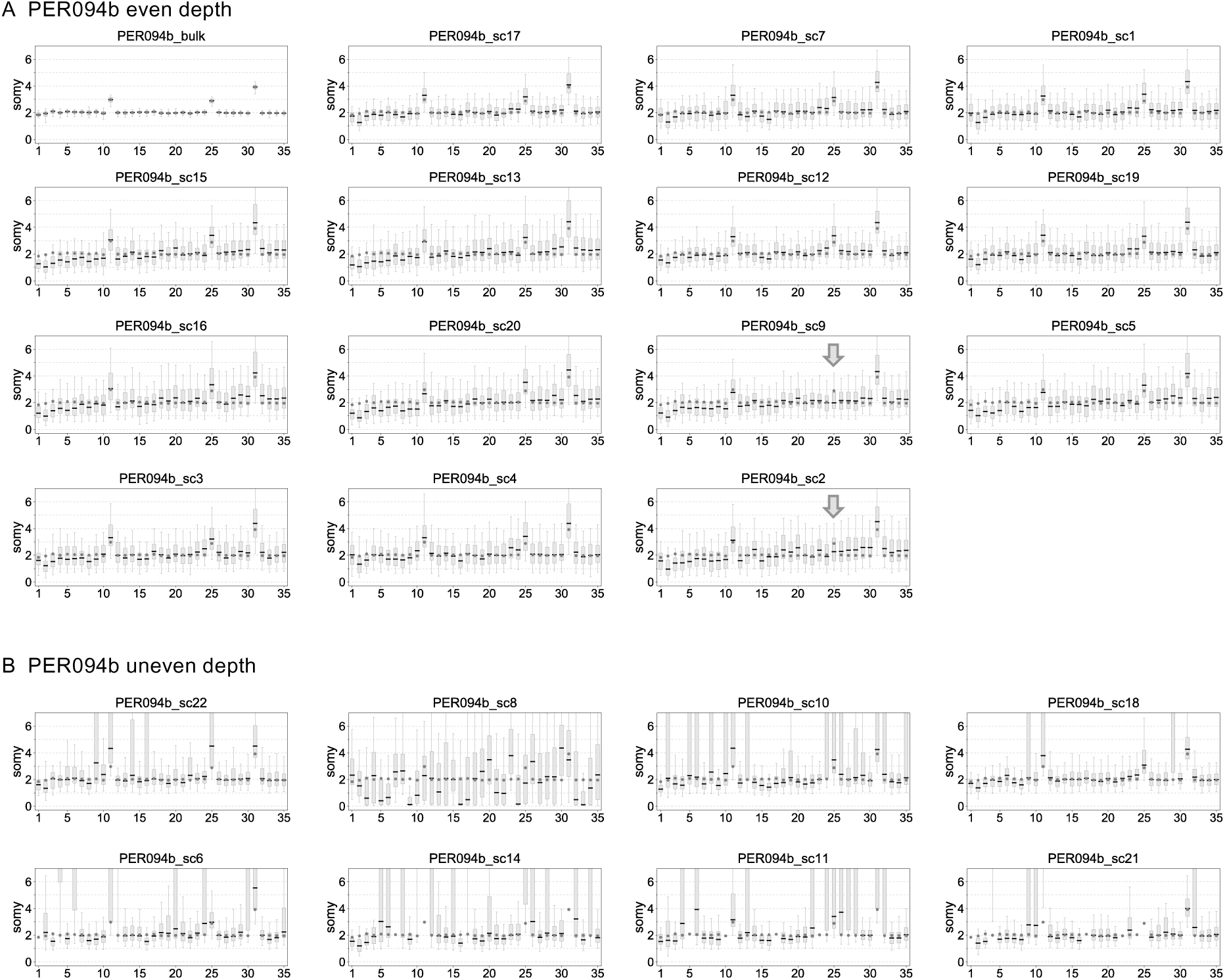
Boxplots of PER094b single cell samples: A) The first boxplot was for the PER094 control. There were 14 even depth single cells that had a short box and short error whiskers, allowing accurate somy estimation. Chromosome 25 is trisomic in most samples, except PER094b_sc9 and sc2, where it is disomic, marked by the arrow. B) The uneven depth of 8 samples led to an erratic somy estimation that could not be corrected. The grey arrows indicate the disomic chromosome 25.

**Supplementary Figure 8.**
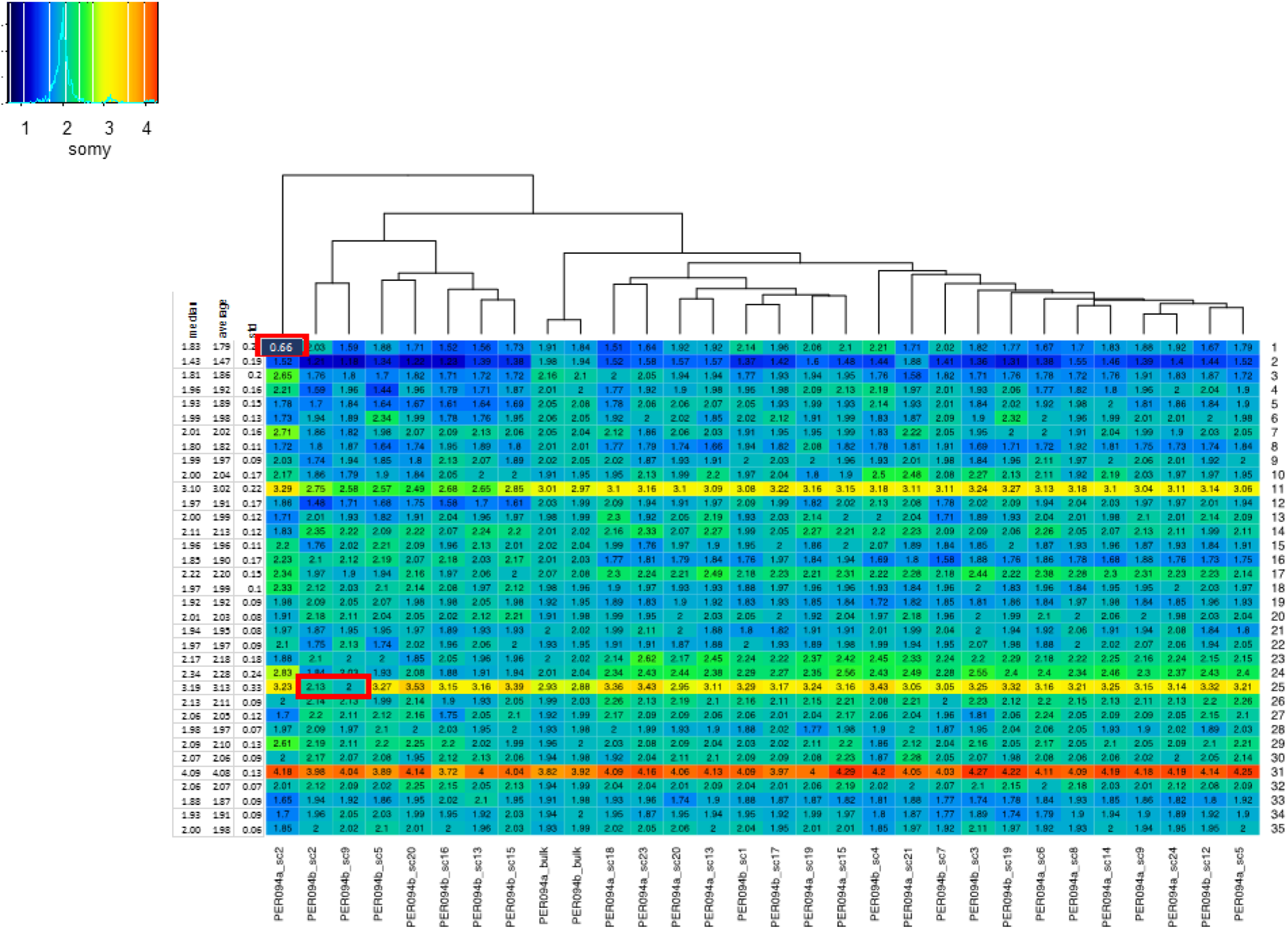
Somy values with GC bias correction for 28 high quality, even depth single cells. The disomic chromosome 25 of PER093b_sc2 and PER093b_sc9 and the monosomic chromosome 1 of PER094a_sc2 are marked in a red box. The color key shows the normalized chromosome read depth, which is corresponding to GC normalized somy.

**Supplementary Figure 9.**
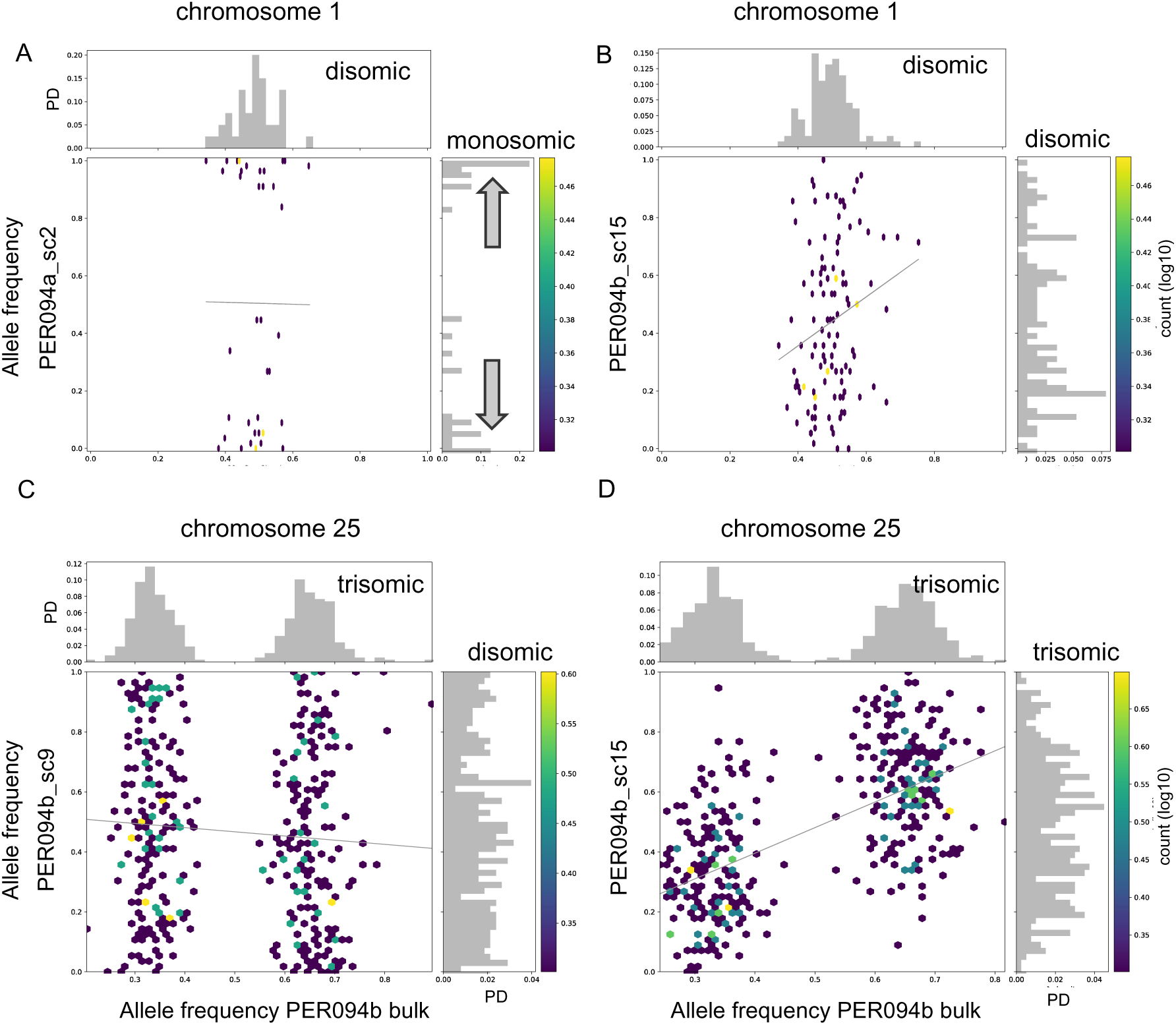
A) Monosomic allele frequency distribution of chromosome 1 of PER094a_sc2 was shown on the y-axis while the clock-like allele frequency distribution of PER094 control (disomic, bulk sequencing) was shown on the x-axis. Two grey arrows indicate the allele frequency shift towards 0 and 1 in PER094a_sc2. B) Disomic chromosome of PER094b_sc13 does not show a clock-like allele frequency distribution, but a flat one, probably because of allele dropouts. C) x-axis shows the bimodal allele frequency distribution of chromosome 25 in PER094 control (trisomic, bulk sequencing); y-axis, ‘flat’ allele frequency distribution in chromosome 25 of PER094b_sc9, shown to be disomic by normalized read-depth. D) y-axis, bimodal distribution of chromosome 25 in PER094b_sc15 shown to be trisomic by normalized depth. The color of dots represents the counts in log10 scale and the straight lines represent the regression lines. PD stands for probability density.

## Supplementary data

**Supplementary_data1.xlsx** file containing 3 tables (S1 to S3)

**Supplementary Table S1.** Mapping statistics and read depth statistics for BPK275 samples were given. The summary table of somy accuracy is also given for comparison.

**Supplementary Table S2.** Mapping statistics and read depth statistics for PER094 samples were given. The summary table of somy accuracy is also given for comparison.

**Supplementary Table S3.** Impact of the median depth and normalized depth standard deviation on the somy accuracy expressed by average somy deviation (ASD). We calculated the correlation values and the p-values between these variables.

**Supplementary_data2.pdf**

Manhattan plots showing the read depth across 36 chromosomes of *L. donovani* BPK275 in (p1, A) the bulk control, (p1, B-G) Picoplex samples derived from different cell numbers and and (p1, H-L and p2, A) RepliG samples derived from different cell numbers. The x- and y-axes represent chromosomal position and average depth per 5000 bp, respectively.

**Supplementary_data3.pdf**

Manhattan plots showing the read depth across 35 chromosomes of *L. braziliensis* PER094a in (p1, A) the bulk control and (p1, B-L; p2, A-L; p3, A-B) Picoplex samples derived from different single cells. Plots are organized from higher to lower quality samples. The x- and y-axes represent chromosomal position and average depth per 5000 bp, respectively.

**Supplementary_data4 file**

Manhattan plots showing the read depth across 35 chromosomes of *L. braziliensis* PER094b in (p1, A) the bulk control and (p1, B-L; p2, A-K) Picoplex samples derived from different single cells. Plots are organized from higher to lower quality samples. The x- and y-axes represent chromosomal position and average depth per 5000 bp, respectively.

**Supplementary_data5.pdf**

GC uncorrected (purple) and GC normalized (green) somy values in *L. braziliensis* PER094a: (p1, A) bulk control and (p1, B-L; p2, A-L; p3, A-B) PicoPlex samples from different single cells. Graphs are organized from higher to lower quality samples. The x- and y-axes represent chromosome numbers and somy, respectively.

**Supplementary_data6.pdf**

GC uncorrected (purple) and GC normalized (green) somy values in *L. braziliensis* PER094b: (p1, A) bulk control and (p1, B-L; p2, A-K) PicoPlex samples from different single cells. Graphs are organized from higher to lower quality samples. The x- and y-axes represent chromosome numbers and somy, respectively.

## Supplementary methods

### qPCR to verify quality of the Picoplex-amplified single cell samples before sequencing

Four qPCR were used, 3 targeting *Leishmania* DNA (kDNA, 18S rDNA and G6PD) and 1 targeting human DNA (RPL30)

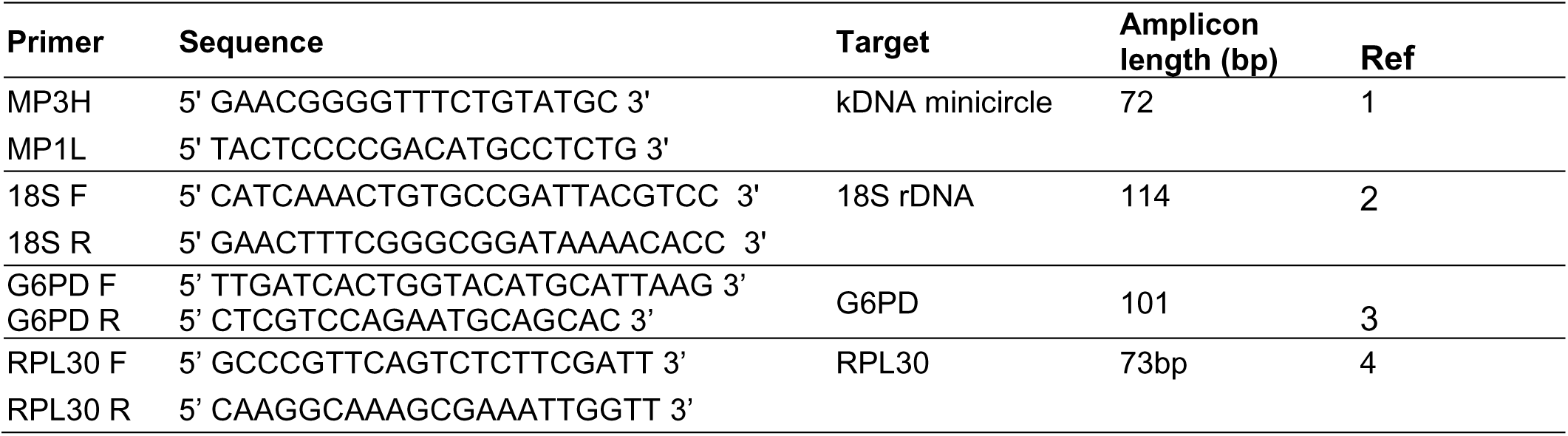

The following cycling conditions were used:

kDNA (*Leishmania braziliensis*): 94°C for 2 mins, followed by 29 rounds of 94°C for 1 min and 54°C for 1 min

18S rDNA (*Leishmania*): 95°C for 3 min, followed by 35 cycles at 95°C for 20 s, 60°C for 20 s and 72°C for 20 s.

G6PD (*Leishmania*): 95°C for 10 mins, followed by 45 rounds of 95°C for 30s, 65°C for 30s, 72°C for 15s.

RPL30 (human): 95°C for 10 mins, followed by 45 rounds of 95°C for 30s, 60°C for 15s, 72°C for 15s.

For the QPCR, we used the Lightcycler 480 (Roche)

To select samples for whole genome sequencing we checked the following parameters :

1. Final concentration of amplified DNA as measured by nanodrop was at least 5 ng/µl
2. The Ct value for the RPL30 was 35 or higher, which is comparable to the values found in pure Leishmania DNA samples.
3. The presence of Leishmania DNA was confirmed by qPCR by at least one of the *Leishmania* markers.
4. Two samples were added based only on the amount of DNA, while the presence of Leishmania DNA was suggestive, but uncertain.

## References

1. Alvar, J. et al. Leishmaniasis worldwide and global estimates of its incidence. PLoS One 7, e35671 (2012).

2. Requena, J. M. Lights and shadows on gene organization and regulation of gene expression in Leishmania. Front. Biosci. (Landmark Ed. 16, 2069–85 (2011).

3. Victoir, K. & Dujardin, J. C. How to succeed in parasitic life without sex? Asking Leishmania. Trends Parasitol. 18, 81–5 (2002).

4. Berg, M., Mannaert, A., Vanaerschot, M., Van Der Auwera, G. & Dujardin, J. C. (Post-) Genomic approaches to tackle drug resistance in Leishmania. Parasitology 140, 1492– 1505 (2013).

5. Imamura, H. et al. Evolutionary genomics of epidemic visceral leishmaniasis in the Indian subcontinent. Elife 5, e12613 (2016).

6. Dumetz, F. et al. Modulation of aneuploidy in leishmania donovani during adaptation to different in vitro and in vivo environments and its impact on gene expression. MBio 8, (2017).

7. Shaw, C. D. et al. In vitro selection of miltefosine resistance in promastigotes of Leishmania donovani from Nepal: Genomic and metabolomic characterization. Mol. Microbiol. 99, 1134–1148 (2016).

8. Sterkers, Y. et al. Novel insights into genome plasticity in Eukaryotes: Mosaic aneuploidy in Leishmania. Molecular Microbiology vol. 86 15–23 (2012).

9. Sterkers, Y., Crobu, L., Lachaud, L., Pagès, M. & Bastien, P. Parasexuality and mosaic aneuploidy in Leishmania: Alternative genetics. Trends in Parasitology vol. 30 429–435 (2014).

10. Macaulay, I. C. & Voet, T. Single Cell Genomics: Advances and Future Perspectives. PLoS Genet. 10, e1004126 (2014).

11. Chen, W. et al. Single Cell Omics: From Assay Design to Biomedical Application. Biotechnol. J. e1900262 (2019) doi:10.1002/biot.201900262.

12. Nair, S. et al. Single-cell genomics for dissection of complex malaria infections. Genome Res. 24, 1028–1038 (2014).

13. Troell, K. et al. Cryptosporidium as a testbed for single cell genome characterization of unicellular eukaryotes. BMC Genomics (2016) doi:10.1186/s12864-016-2815-y.

14. Zhang, C. Z. et al. Calibrating genomic and allelic coverage bias in single-cell sequencing. Nat. Commun. 6, 6822 (2015).

15. Chen, D. Y. et al. Comparison of single cell sequencing data between two whole genome amplification methods on two sequencing platforms. Sci. Rep. 8, 4963 (2018).

16. Garvin, T. et al. Interactive analysis and assessment of single-cell copy-number variations. Nature Methods vol. 12 1058–1060 (2015).

17. Rogers, M. B. et al. Chromosome and gene copy number variation allow major structural change between species and strains of Leishmania. Genome Res. 21, 2129–2142 (2011).

18. Domagalska, M. A. et al. Genomes of Leishmania parasites directly sequenced from patients with visceral leishmaniasis in the Indian subcontinent. PLoS Negl. Trop. Dis. 13, e0007900 (2019).

19. Barja, P. P. et al. Haplotype selection as an adaptive mechanism in the protozoan pathogen Leishmania donovani. Nat. Ecol. Evol. 1, 1961–1969 (2017).

20. Downing, T. et al. Whole genome sequencing of multiple Leishmania donovani clinical isolates provides insights into population structure and mechanisms of drug resistance. Genome Res. 21, 2143–56 (2011).

21. Domagalska, M. A. et al. Genomes of intracellular Leishmania parasites directly sequenced from patients. bioRxiv 676163 (2019) doi:10.1101/676163.

22. de Bourcy, C. F. A. et al. A quantitative comparison of single-cell whole genome amplification methods. PLoS One 9, e105585 (2014).

23. Barea, J. S., Lee, J. & Kang, D. K. Recent advances in droplet-based microfluidic technologies for biochemistry and molecular biology. Micromachines vol. 10 (2019).

24. Jara, M. et al. Tracking of quiescence in Leishmania by quantifying the expression of GFP in the ribosomal DNA locus. bioRxiv 641290 (2019) doi:10.1101/641290.

25. Kozarewa, I. et al. Amplification-free Illumina sequencing-library preparation facilitates improved mapping and assembly of (G+C)-biased genomes. Nat. Methods 6, 291–295 (2009).

26. Bronner, I. F., Quail, M. A., Turner, D. J. & Swerdlow, H. Improved Protocols for Illumina Sequencing. Curr. Protoc. Hum. Genet. 80, 18.2.1–42 (2014).

27. Deleye, L. et al. Performance of four modern whole genome amplification methods for copy number variant detection in single cells. Sci. Rep. 7, (2017).

28. McKenna, A. et al. The genome analysis toolkit: A MapReduce framework for analyzing next-generation DNA sequencing data. Genome Res. 20, 1297–1303 (2010).

29. Dumetz, F. et al. Modulation of aneuploidy in leishmania donovani during adaptation to different in vitro and in vivo environments and its impact on gene expression. MBio 8, (2017).

30. Benjamini, Y. & Speed, T. P. Summarizing and correcting the GC content bias in high-throughput sequencing. Nucleic Acids Res. (2012) doi:10.1093/nar/gks001.

31. Scheinin, I. et al. DNA copy number analysis of fresh and formalin-fixed specimens by shallow whole-genome sequencing with identification and exclusion of problematic regions in the genome assembly. Genome Res. (2014) doi:10.1101/gr.175141.114.

32. Williams, T., Kelley, C. & many others. Gnuplot 5.2: an interactive plotting program. (2018).

33. Hunter, J. D. Matplotlib: A 2D graphics environment. Comput. Sci. Eng. 9, 90–95 (2007).

## References

1 Lopez, M. et al. Diagnosis of *Leishmania* using the polymerase chain reaction: a simplified procedure for field work. Am. J. Trop. Med. Hyg. 49, 348–56 (1993).

2 Jara, M. et al. Macromolecular biosynthetic parameters and metabolic profile in different life stages of *Leishmania braziliensis*: Amastigotes as a functionally less active stage. PLoS One 12, e0180532 (2017).

3 Castilho, T. M., Camargo, L. M. A., McMahon-Pratt, D., Shaw, J. J. & Floeter-Winter, L. M. A real-time polymerase chain reaction assay for the identification and quantification of American Leishmania species on the basis of glucose-6-phosphate dehydrogenase. Am. J. Trop. Med. Hyg. 78, 122–132 (2008).

4 Domagalska, M. A., et al. Genomes of Leishmania parasites directly sequenced from patients with visceral leishmaniasis in the Indian subcontinent. PLoS Negl. Trop. Dis. 13, e0007900 (2019).

