## Supplementary data 2 for "Evaluation of whole genome amplification and bioinformatic methods for the characterization of *Leishmania* genomes at a single cell level"

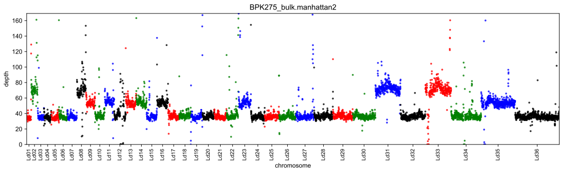


(A) BPK275_bulk

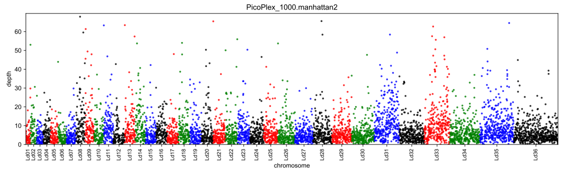


(B) PicoPlex_1000

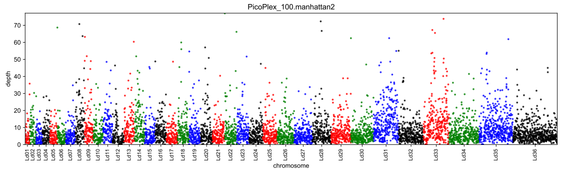


(C) PicoPlex_100

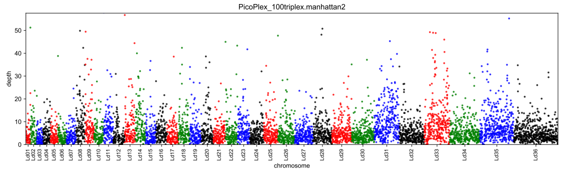


(D) PicoPlex_100triplex

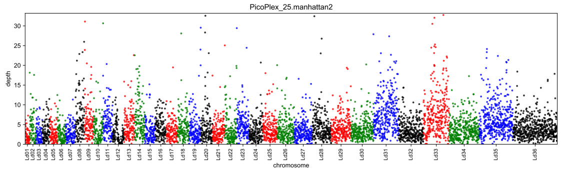


(E) PicoPlex_25

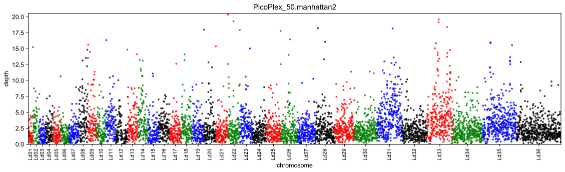


(F) PicoPlex_50

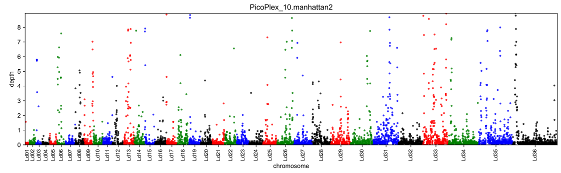


(G) PicoPlex_10

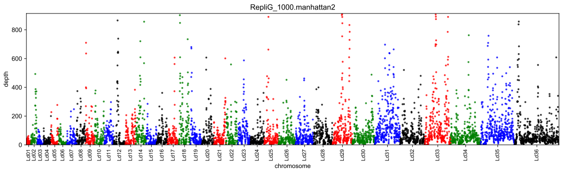


(H) RepliG_1000

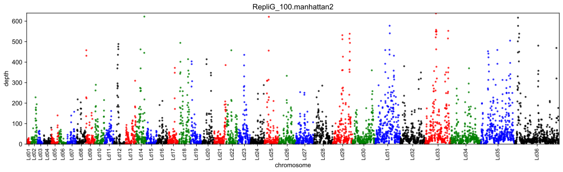


(I) RepliG_100

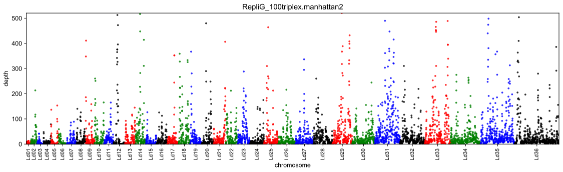


(J) RepliG_100triplex

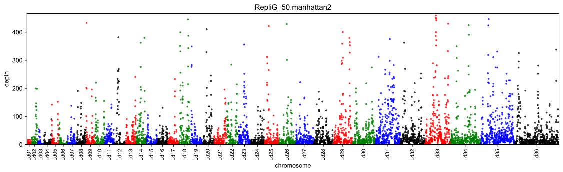


(K) RepliG_50

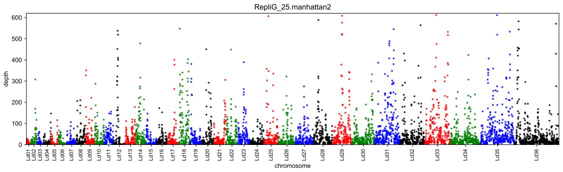


(L) RepliG_25

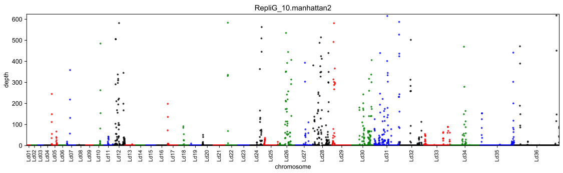
RepliG_10.manhattan2
depth
0
100
200
300
400
500
600
Ld01 Ld02 Ld03 Ld04 Ld05 Ld06 Ld07 Ld08 Ld09 Ld10 Ld11 Ld12 Ld13 Ld14 Ld15 Ld16 Ld17 Ld18 Ld19 Ld20 Ld21 Ld22 Ld23 Ld24 Ld25 Ld26 Ld27 Ld28 Ld29 Ld30 Ld31 Ld32 Ld33 Ld34 Ld35 Ld36
chromosome

(A) RepliG_10
