## Supplementary data 3 for "Evaluation of whole genome amplification and bioinformatic methods for the characterization of *Leishmania* genomes at a single cell level"

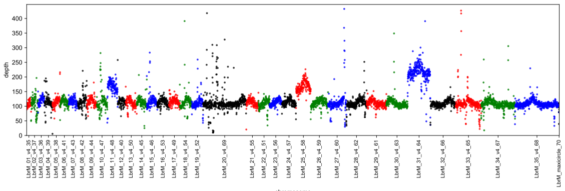


(A) PER094a_bulk

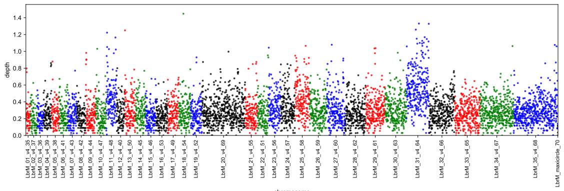


(B) PER094a_sc19

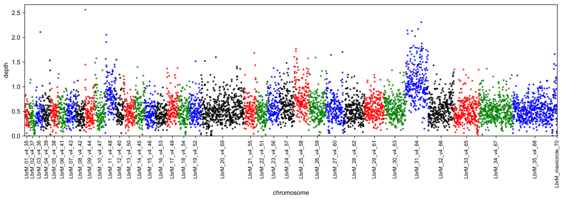


(C) PER094a_sc20

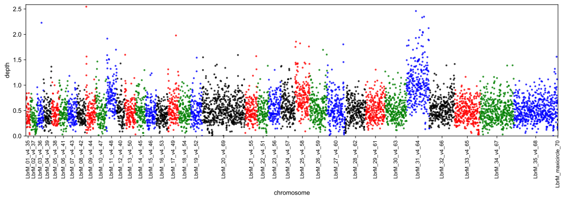


(D) PER094a_sc8

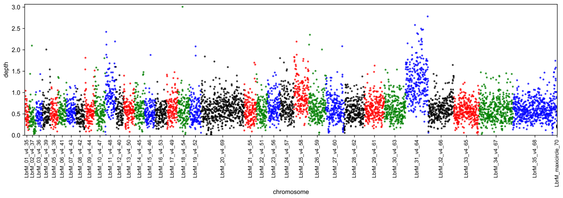


(E) PER094a_sc9

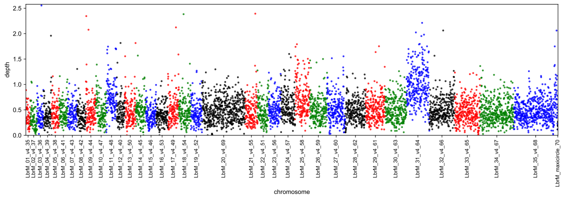


(F) PER094a_sc6

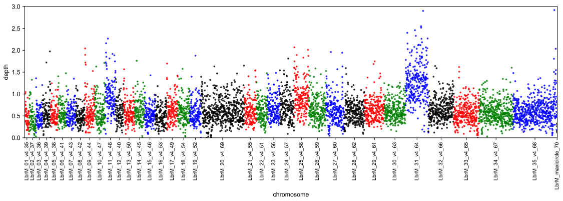


(G) PER094a_sc24

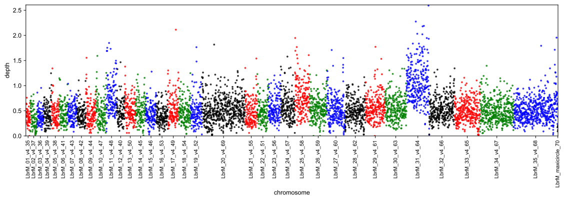


(H) PER094a_sc5

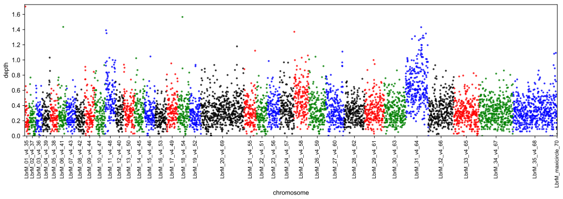


(I) PER094a_sc18

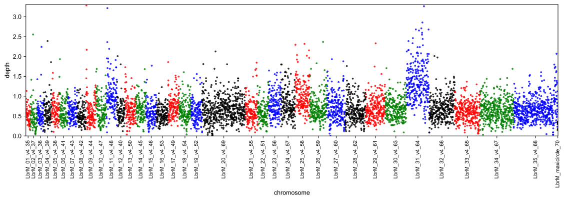


(J) PER094a_sc13

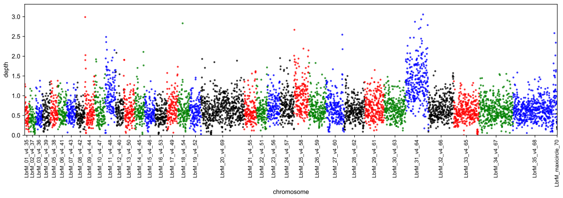


(K) PER094a_sc14

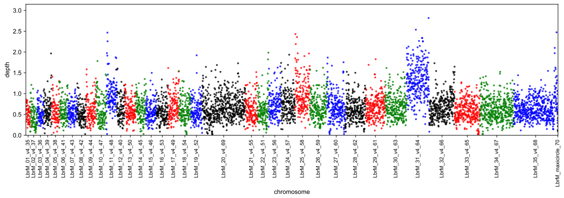


(L) PER094a_sc15

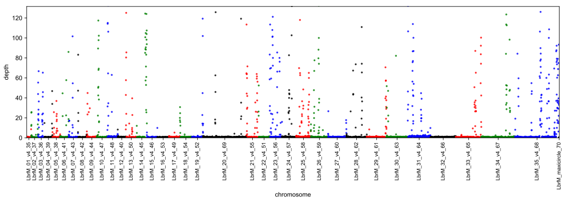


(A) PER094a_sc23

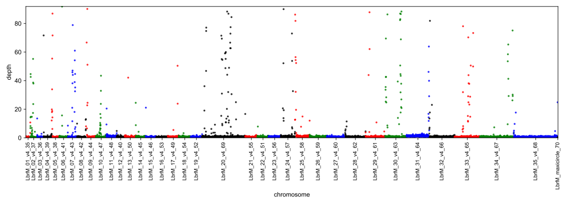


(B) PER094a_sc21

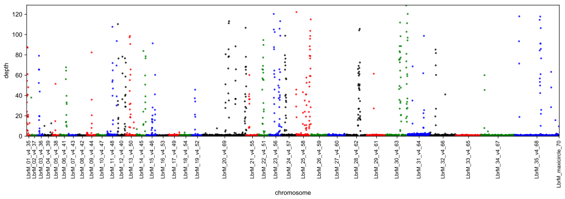


(C) PER094a_sc7

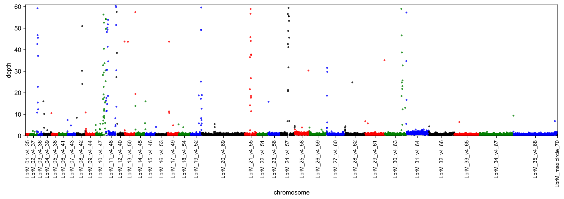


(D) PER094a_sc11

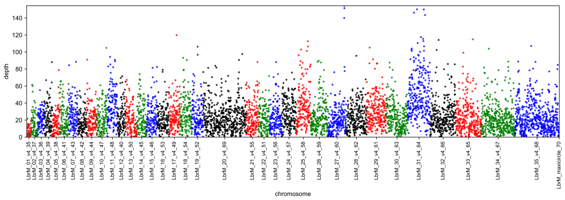


(E) PER094a_sc2

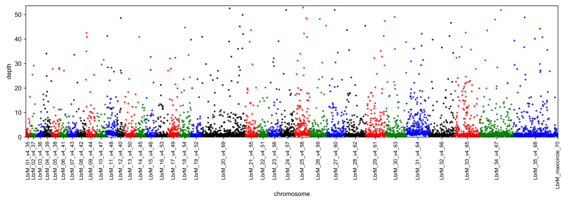


(F) PER094a_sc10

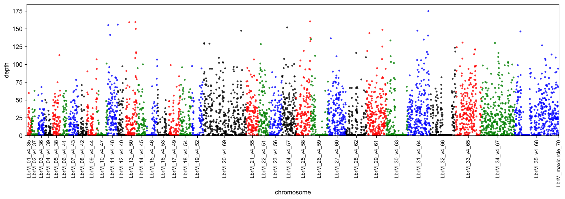


(G) PER094a_sc17

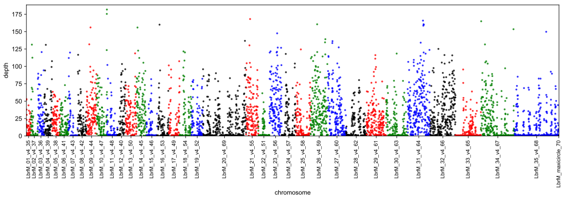


(H) PER094a_sc16

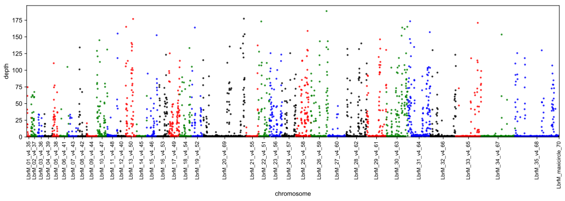


(I) PER094a_sc12

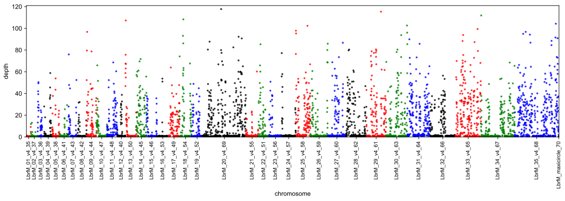


(J) PER094a_sc25

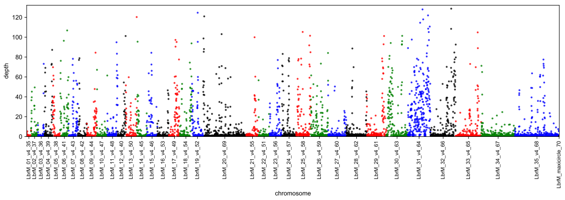


(K) PER094a_sc3

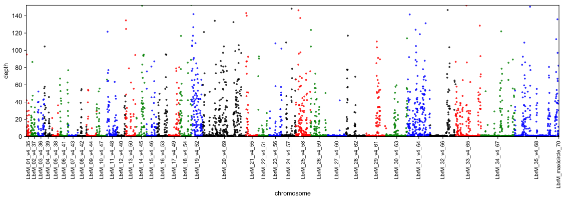


(L) PER094a_sc1

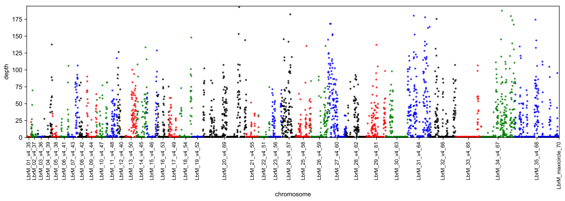


(A) PER094a_sc22

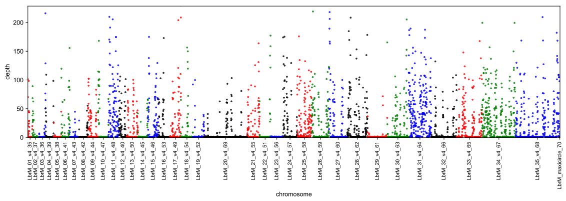


(B) PER094a_sc4
