## Supplementary data 4 for "Evaluation of whole genome amplification and bioinformatic methods for the characterization of *Leishmania* genomes at a single cell level"

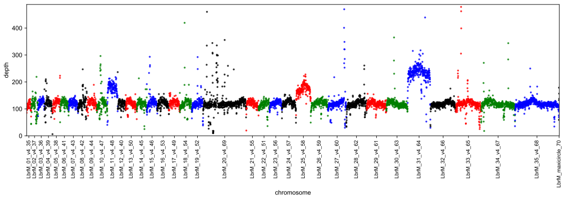

(A) PER094b_bulk

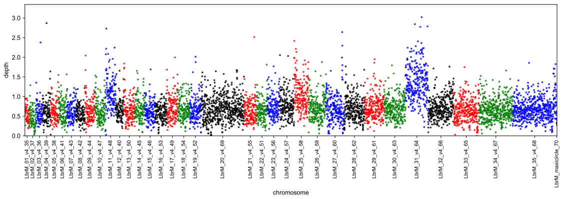

(B) PER094b_sc17

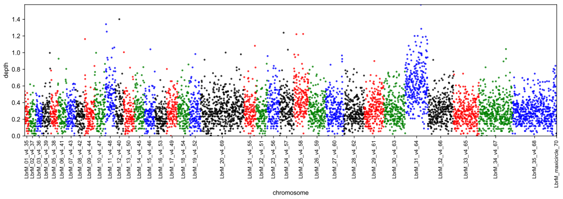

(C) PER094b_sc7

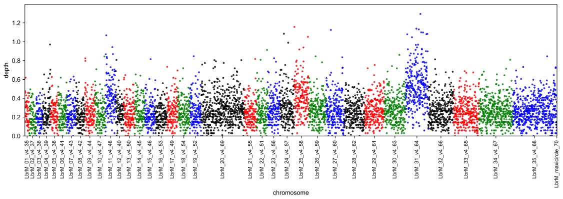

(D) PER094b_sc1

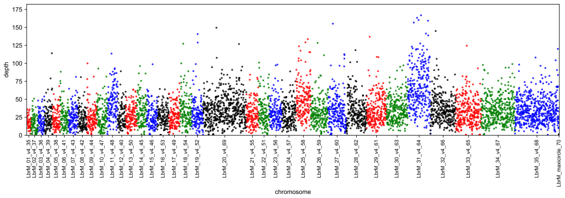

(E) PER094b_sc15

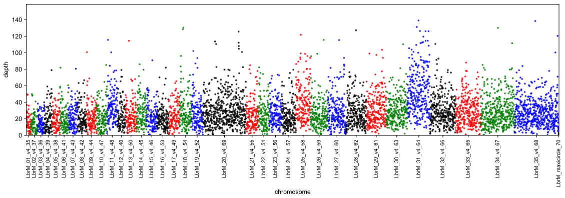

(F) PER094b_sc13

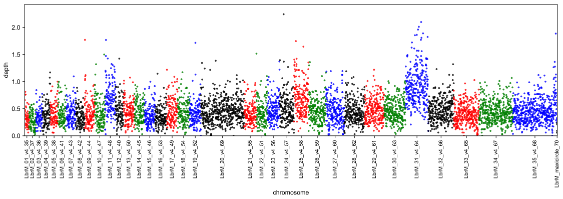

(G) PER094b_sc12

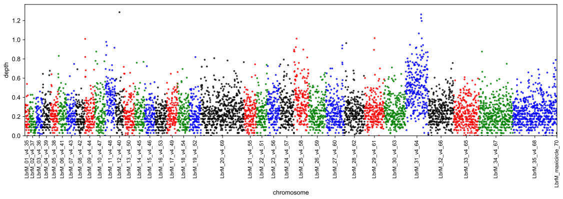

(H) PER094b_sc19

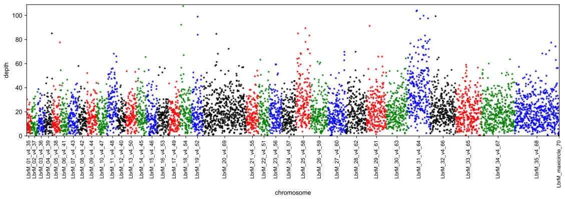

(I) PER094b_sc16

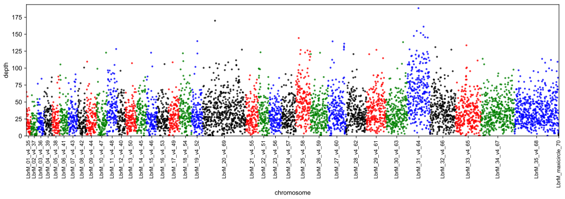

(J) PER094b_sc20

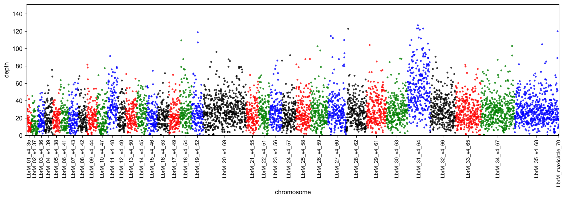

(K) PER094b_sc9

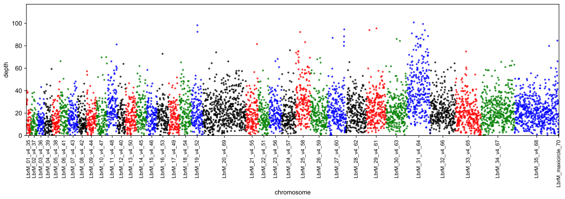

(L) PER094b_sc5

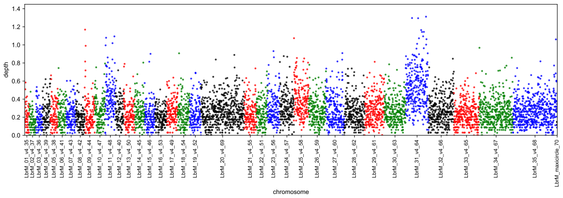

(A) PER094b_sc3

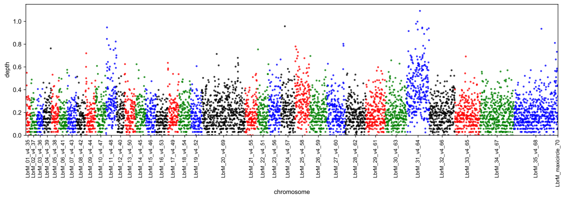

(B) PER094b_sc4

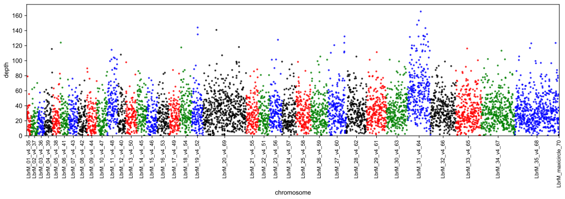

(C) PER094b_sc2

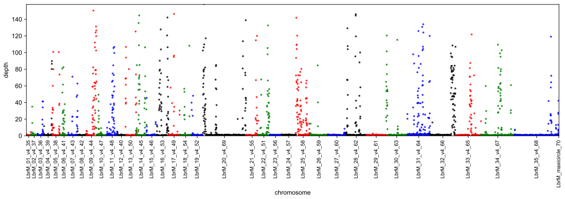

(D) PER094b_sc22

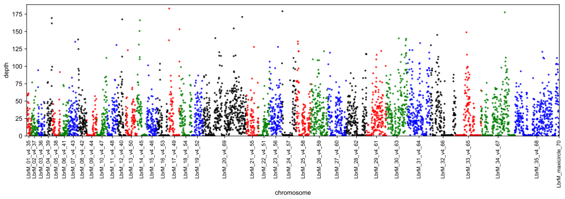

(E) PER094b_sc8

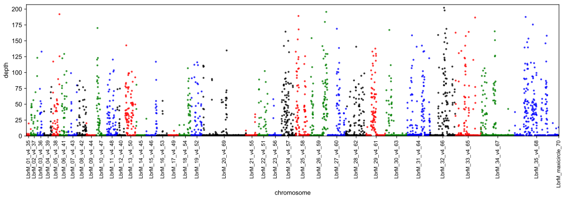

(F) PER094b_sc10

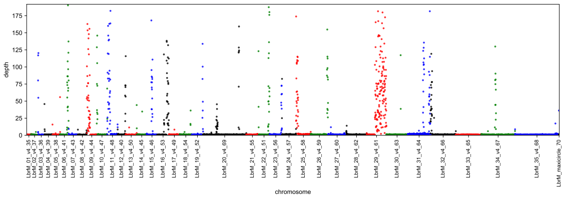

(G) PER094b_sc18

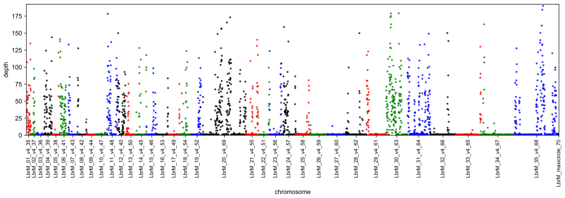

(H) PER094b_sc6

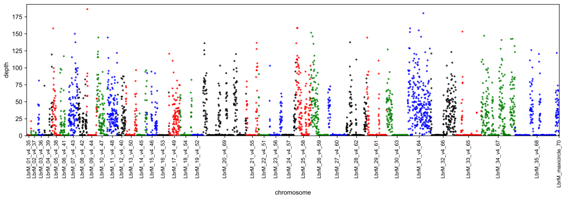

(I) PER094b_sc14

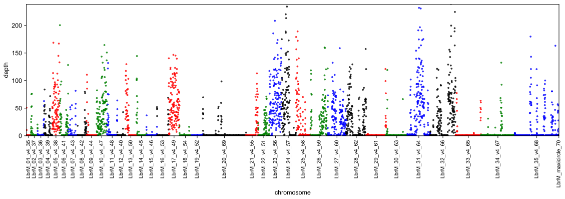

(J) PER094b_sc11

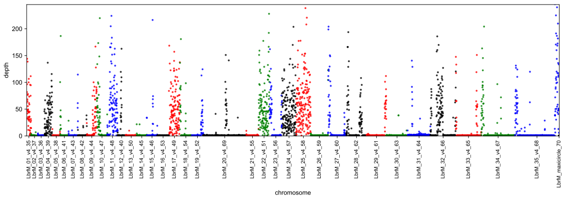

(K) PER094b_sc21
