## Supplementary data 5 for "Evaluation of whole genome amplification and bioinformatic methods for the characterization of *Leishmania* genomes at a single cell level"

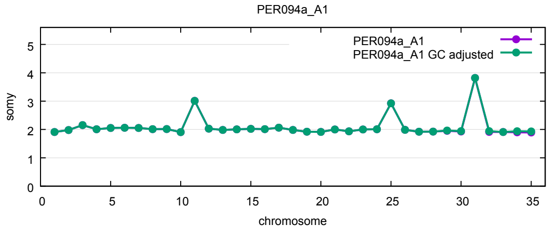

(A) PER094a_bulk

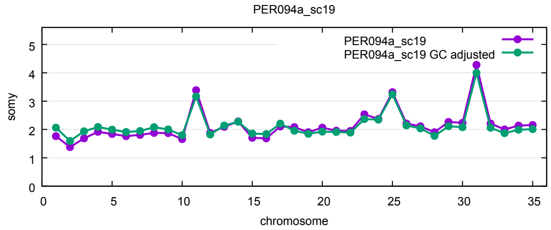

(B) PER094a_sc19

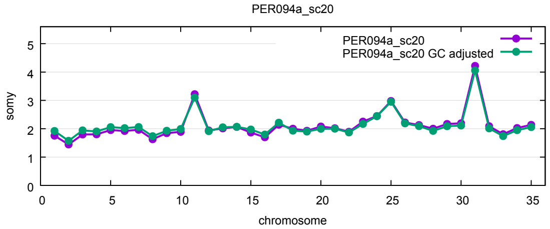

(C) PER094a_sc20

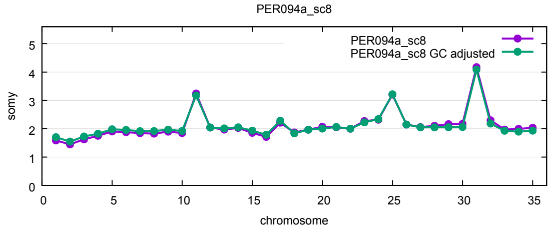

(D) PER094a_sc8

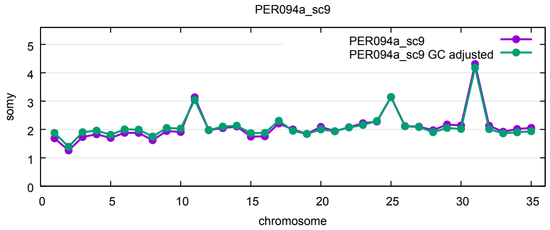

(E) PER094a_sc9

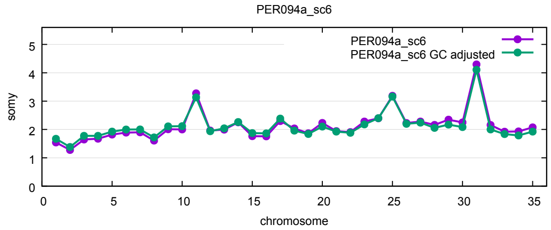

(F) PER094a_sc6

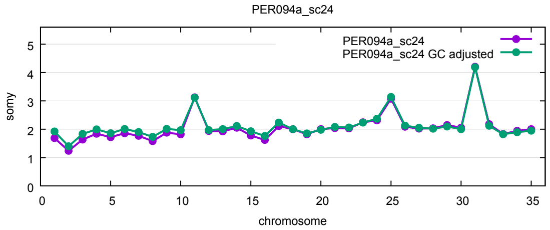

(G) PER094a_sc24

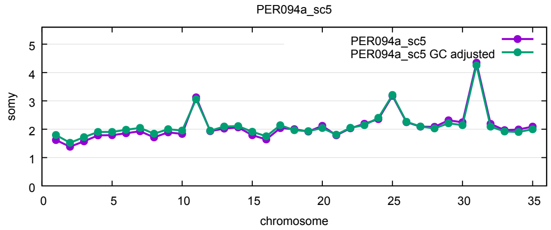

(H) PER094a_sc5

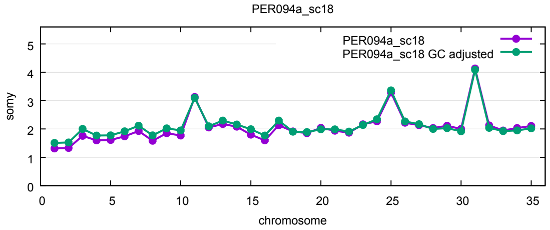

(I) PER094a_sc18

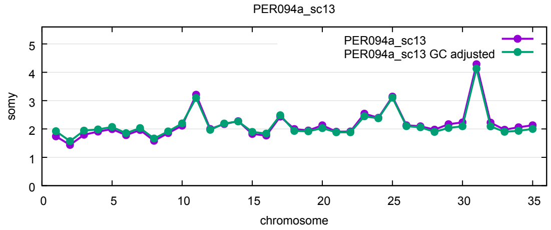

(J) PER094a_sc13

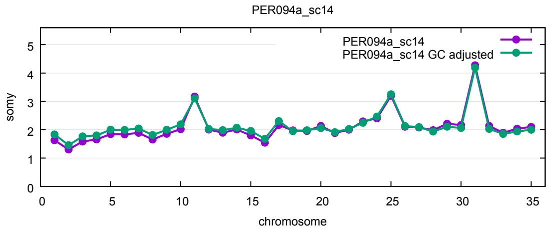

(K) PER094a_sc14

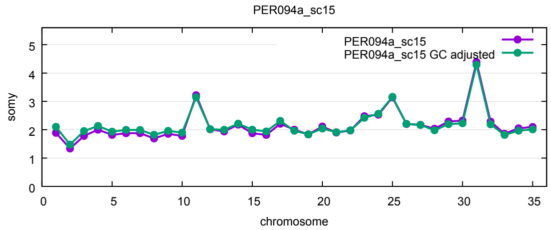

(L) PER094a_sc15

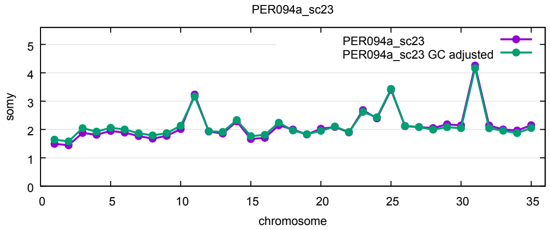

(A) PER094a_sc23

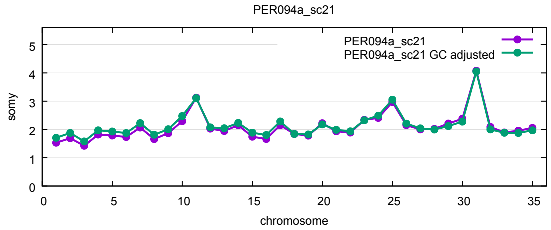

(B) PER094a_sc21

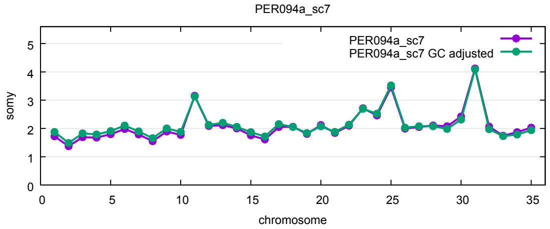

(C) PER094a_sc7

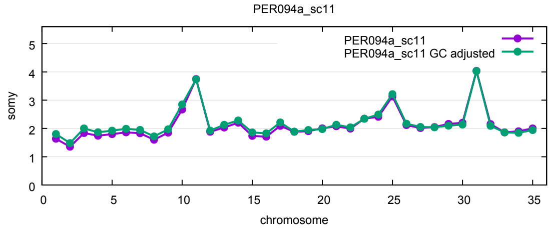

(D) PER094a_sc11

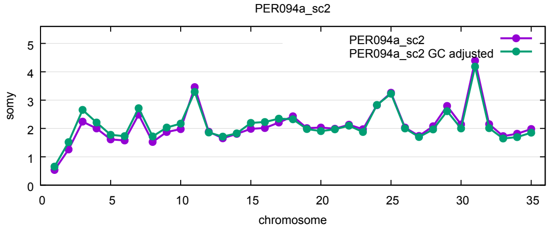

(E) PER094a_sc2

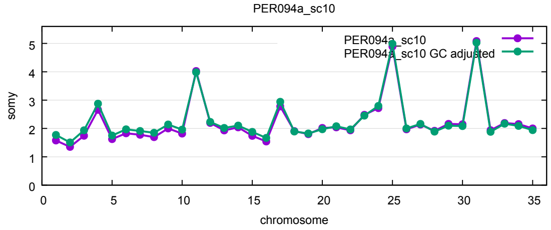

(F) PER094a_sc10

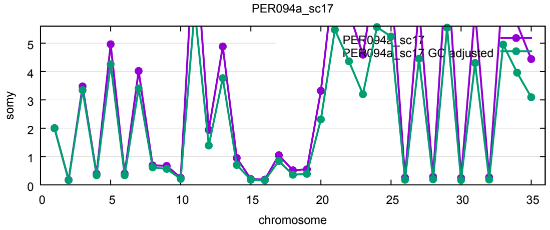

(G) PER094a_sc17

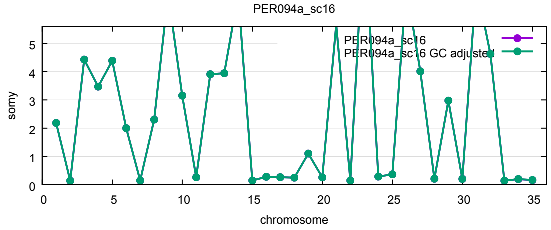

(H) PER094a_sc16

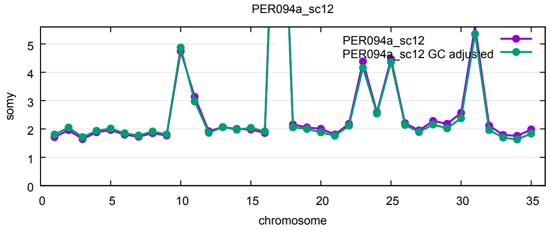

(I) PER094a_sc12

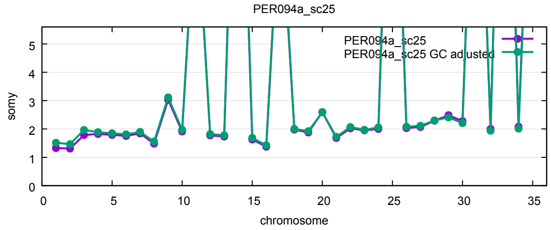

(J) PER094a_sc25

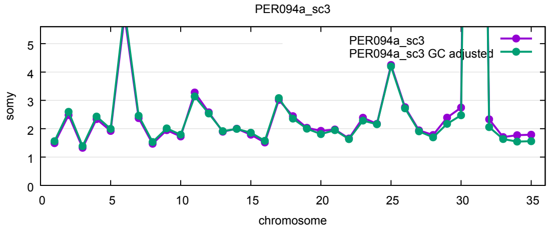

(K) PER094a_sc3

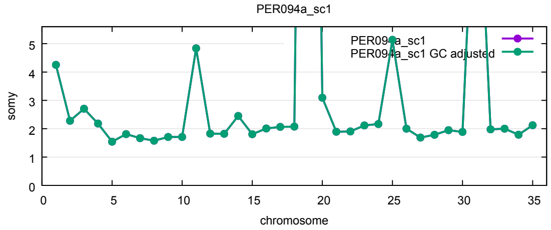

(L) PER094a_sc1

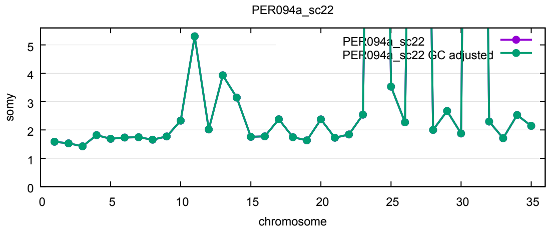


(A) PER094a_sc22

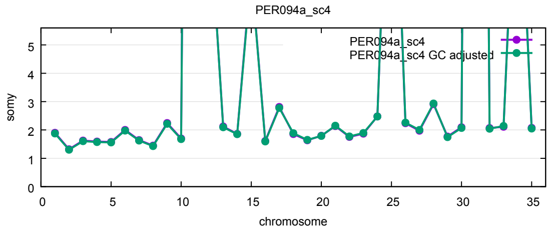


(B) PER094a_sc4
