## Supplementary data 6 for "Evaluation of whole genome amplification and bioinformatic methods for the characterization of *Leishmania* genomes at a single cell level"

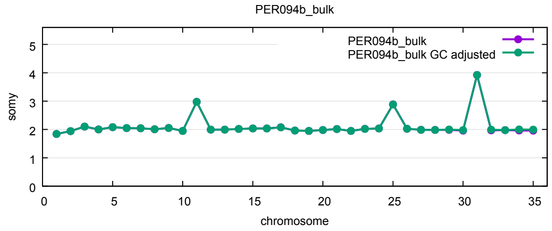


(A) PER094b_bulk

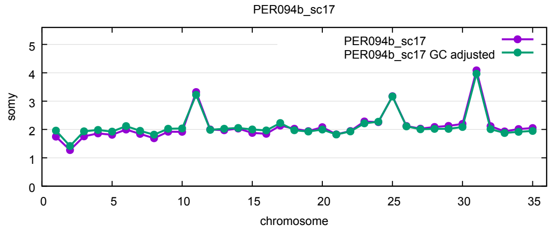


(B) PER094b_sc17

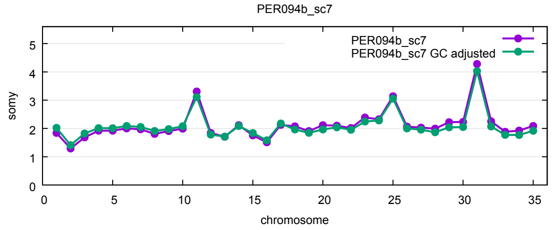


(C) PER094b_sc7

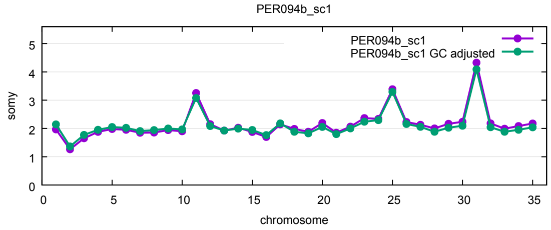


(D) PER094b_sc1

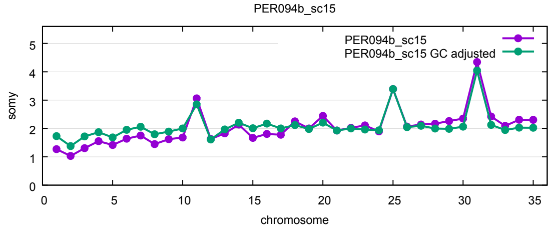


(E) PER094b_sc15

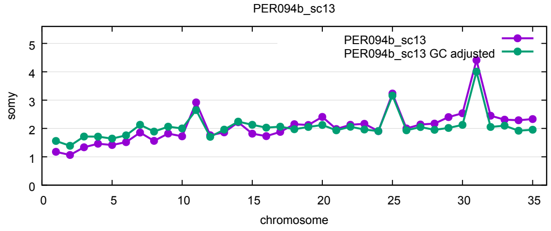


(F) PER094b_sc13

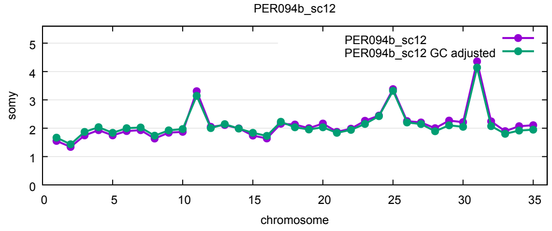


(G) PER094b_sc12

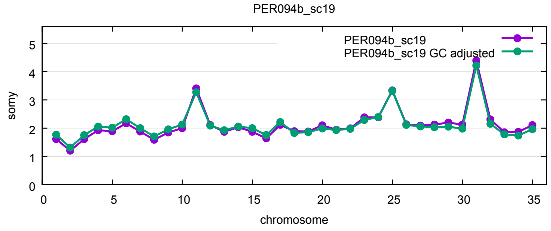


(H) PER094b_sc19

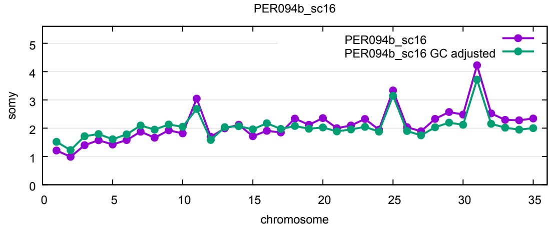


(I) PER094b_sc16

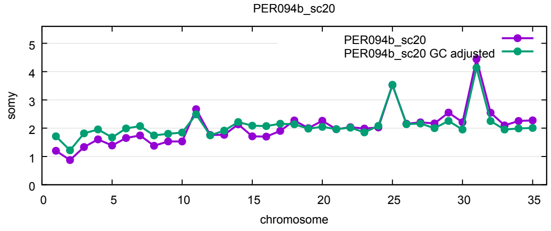


(J) PER094b_sc20

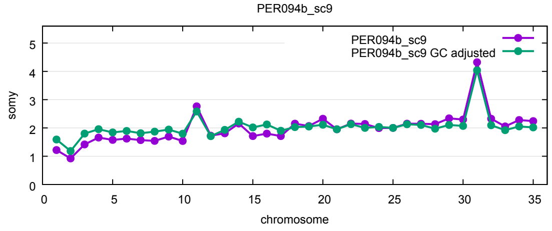


(K) PER094b_sc9

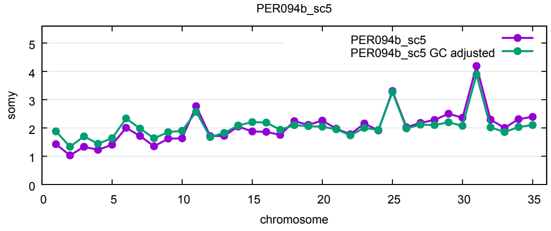


(L) PER094b_sc5

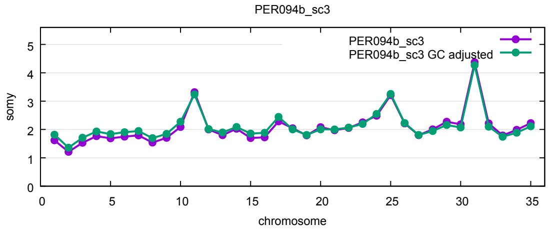

(A) PER094b_sc3

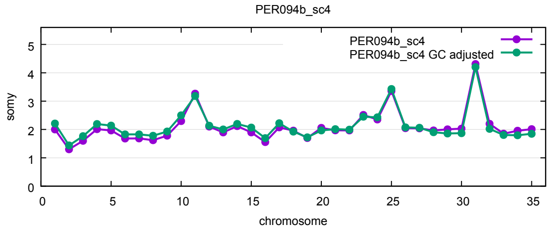

(B) PER094b_sc4

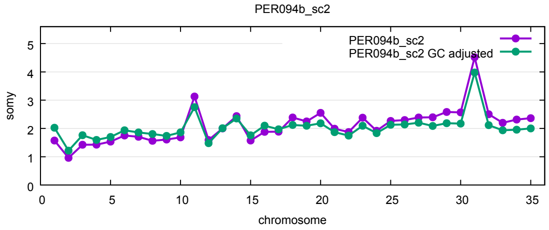

(C) PER094b_sc2

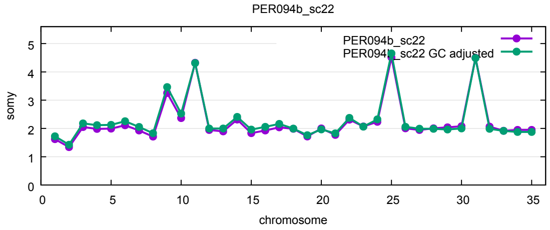

(D) PER094b_sc22

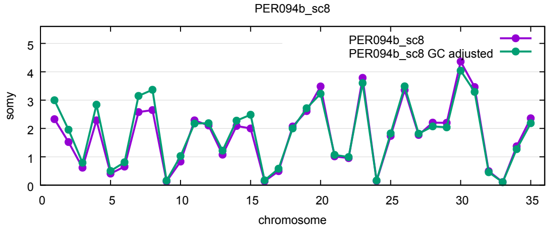

(E) PER094b_sc8

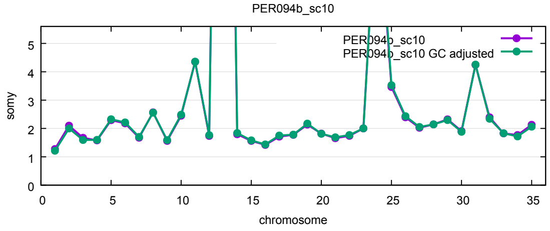

(F) PER094b_sc10

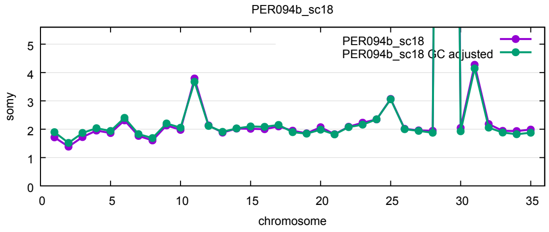

(G) PER094b_sc18

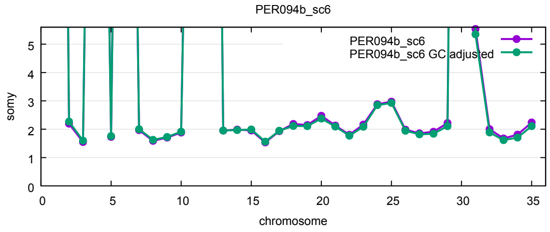

(H) PER094b_sc6

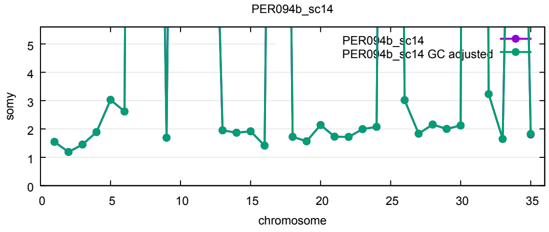

(I) PER094b_sc14

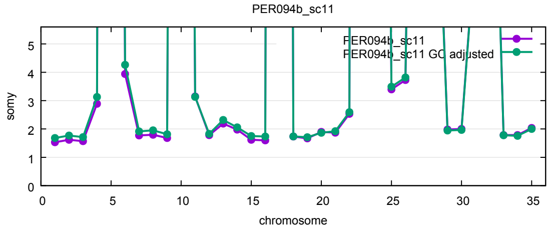

(J) PER094b_sc11

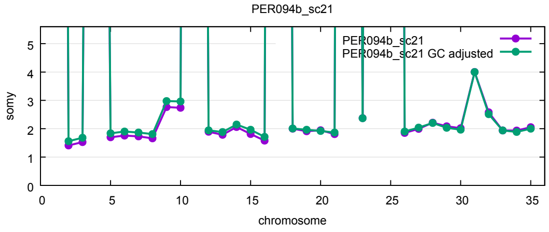

(K) PER094b_sc21
